# Dengue NS1 Antibodies drive Immune Complex Formation, Hyperglycaemia and systemic pathology in a murine NS1 plasmid challenge model

**DOI:** 10.1101/2025.09.25.678465

**Authors:** Chiroshri Dutta, Debrupa Dutta, Soumi Sukla, Subhajit Biswas

## Abstract

Dengue virus (DV) NS1, a secreted virotoxin and key pathogenic factor, can trigger immune responses with poorly understood long-term effects. This study assessed immunopathology in mice administered with DV NS1 plasmid DNA via intraperitoneal (IP), intramuscular (IM), or intravenous (IV) route for DV serotypes 1–4. IP delivery caused the most pronounced effects, including elevated AST/ALT and GRP78 levels, hyperglycemia, and altered organ weights, with DV4 NS1 showing the strongest hepatic damage. Despite serum NS1 antigen being undetectable, mice developed strong NS1-specific antibodies (Abs) and immune complexes. Liver histology revealed degeneration and immune cell depletion. DV NS1 plasmid DNA was detected in liver tissue, but not RNA. DV could infect and replicate in murine pancreatic beta cells. In liver cells, DV increased GAPDH expression, while NS1-Ab-positive serums reduced it. Findings indicated that NS1-specific Abs, not the antigen, drove immune-metabolic dysfunctions, emphasizing the need to evaluate Ab-mediated effects in dengue pathogenesis.

**Significance Statement:** Dengue virus (DV) remains a leading tropical pathogen, yet the long-term effects of its secreted virotoxin NS1 are incompletely understood. Using a murine model, we demonstrated that NS1 plasmid DNA administration drove systemic and organ-specific pathology. Serotype-specific differences in immune responses, biochemical alterations, and histopathological changes underscored NS1’s complex role in disease severity. Elevated immune complexes and liver enzyme profiles highlighted mechanisms of immune modulation and hepatic injury, key features of dengue, while increased serum glucose and GRP78 levels pointed to early markers of diabetes onset. These findings provide foundational evidence that NS1- and NS1 antibody–mediated pathways link dengue pathogenesis with metabolic dysfunction, offering critical insights into host–pathogen interactions and comorbidity development.

## Introduction

Dengue virus (DV) is a single-stranded positive sense RNA virus belonging to family *Flaviviridae.* Dengue, caused by DV, is manifested by febrile condition accompanied by thrombocytopenia and rash mainly in tropical and subtropical countries. These conditions may progress to the perilous dengue haemorrhagic fever and dengue shock syndrome that can eventually lead to multiple organ failure and death.(1, 2) Dengue infection can be caused by four antigenically distinct but genetically similar serotypes of the virus, namely DV1, DV2, DV3 and DV4.(3) Dengue has become a global health threat and severe economic burden with ten times increase in infection rate in the last two decades.(4)

Till date, there is no specific antiviral approved for the treatment of dengue fever. The available treatment regimen includes supportive therapy and in severe cases, platelet transfusion is recommended.(5) One major hurdle behind the understanding of DV pathogenesis and development of dengue therapeutics and vaccines is the lack of an ideal murine model that recapitulates dengue infection in humans.(6) Currently, there are two approved vaccines available for the prevention of dengue namely Dengvaxia and Qdenga. Dengvaxia has limited use and can be given to 9-40 years old seropositive individuals, and it carries the potential risk of developing antibody (Ab)-dependent enhancement (ADE) of infection whereas Qdenga has shown only moderate efficacy for serotypes other than DV2.(7, 8)

Dengue virus consists of three structural (C, prM and E) and seven non-structural proteins (NS1, NS2a, NS2b, NS3, NS4a, NS4b and NS5).(9) Among these, NS1 is the most abundant antigenic non-structural protein (46-55 kDa) primarily responsible for the pathogenesis of dengue and is an important diagnostic marker.(10, 11) NS1 plays a role in virus replication and gets transported to cell surface where it is associated with the cell membrane and secreted as a soluble hexamer.(12) DV NS1 has the potential to disrupt the integrity of endothelial cells due to the activation of macrophages eliciting production of inflammatory cytokines.(13) NS1-mediated disruption of endothelial glycocalyx layer increases endothelial permeability causing vascular leakage in human pulmonary microvascular endothelial cells.(14) The dual role of DV NS1 in virus replication and pathogenesis makes it an attractive antiviral target, as blocking NS1 with Abs, vaccines, or small molecules can simultaneously impair viral RNA replication and neutralize NS1-mediated damage without causing ADE.(15)

Highly immunogenic nature of NS1 makes it a potential vaccine candidate as it generates strong humoral response in dengue infected individuals. The pathogenic responses of NS1 have been shown to be blocked by monoclonal antibodies (mAb) to NS1.(16) However, NS1-mediated pathogenesis has been shown to be strain-dependent. So, it is a challenge to develop it as a universal DV immunogen.(17) Previous reports suggested that immunocompetent mice when immunized with a DNA vaccine coding for NS proteins, showed T-cell mediated protective response against dengue infection.(18)

In the present study, we aimed to demonstrate the impact of expressing DV NS1 of different serotypes in mice using different routes of administration such as intraperitoneal (IP), intramuscular (IM) and intravenous (IV). We have also studied the NS1 immune response, and the impact of NS1 Abs (immunopathology) in the murine model.

## Results

### DV NS1 plasmid administration altered survival, organ weights, and liver enzyme profiles

Animal experiments were performed in two sets. In the first set, 20□µg DV2 NS1 plasmid was administered via IP route with or without FuGENE, and also by IM and IV routes without FuGENE. All animals in this set survived till the end of the experiment, yielding a 100% survival rate. Based on these findings, a second set was designed using the IP route and higher plasmid dose (100µg/animal), to assess serotype-specific effects with DV1–DV4 NS1. In this phase, a mouse in the DV2-NS1 group died one week post-first dose, resulting in 80% survival for this group. Two mice in the DV4-NS1 group died, one after the first dose by the second week, and another following the third dose by the sixth week, reducing survival rate to 60%. All remaining mice in DV1, DV3, and control groups survived (**Figure 1A**). Overall, the fatality rate among NS1-inoculated mice (n=48) was 6.25%, with three deaths observed, while all control mice (n=15) survived.

**Figure 1:**
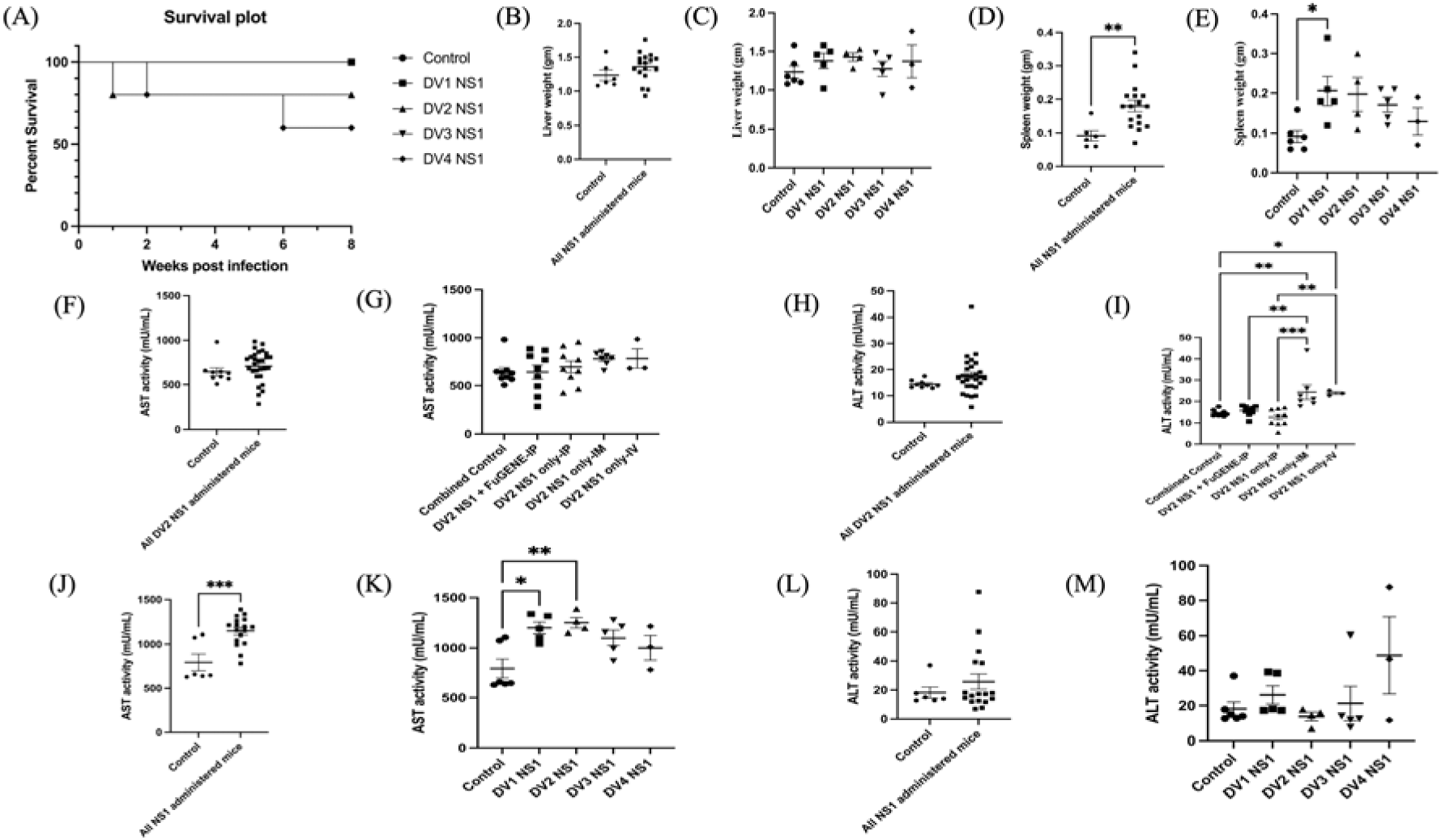
Effects of dengue NS1 inoculation in mice on survival, organ weights, and liver enzyme levels. (**A**) Survival plot of serotype specific DV NS1 administered mice. One animal from the DV2 NS1 group died within the 1^st^ week, and two animals from the DV4 NS1 group died during the 2^nd^ and 6^th^ weeks, respectively. (**B–C**) Combined and individual serotype-specific liver weights in DV NS1 administered mice compared to controls. (**D–E**) Combined and individual serotype-specific spleen weights in the same groups. (**F–I**) Serum AST and ALT levels in DV2 NS1 plasmid-inoculated mice via IP, IM, and IV routes. (F,H) Mean AST and ALT levels in DV2 NS1 plasmid inoculated mice (all routes combined) versus controls. (G,I) Route-specific AST and ALT levels for DV2 NS1 plasmid administered mice compared to combined controls (control and FuGENE control groups). (**J–M**) Serum AST and ALT levels in mice inoculated with NS1 plasmid of four different DV serotypes via IP route. (J,L) Mean AST and ALT levels in mice administered with serotype specific DV NS1 plasmids combined compared to control. (K,M) Serotype-specific AST and ALT levels in mice administered with NS1 plasmids via IP route. Data represented as mean±SEM. *p<0.05; **p<0.01; ***p<0.001. D=Days post DV NS1 administration.

Body weights remained largely unchanged across treatment groups, except for an increase in DV2 NS1 IV-injected mice at 7 and 14 dpi (**Figure S1A**). No significant changes were noted in DV2 NS1 IP or IM groups, or in mice receiving DV1–4 NS1 via IP route (**Figures S1A and S1B**). Organ weights (liver and spleen) were recorded after sacrificing the animals (**Figures 1B-E**). In set 2 experiments, no significant differences were observed in the liver weights (**Figures 1B and 1C**), although a little rise was observed in the mean liver weight of DV1-4 NS1 groups combined compared to the controls (**Figure 1B)**. In contrast, spleen weights were significantly higher in all NS1-immunized mice combined (**Figure 1D**), with DV1-NS1 inducing a significant increase relative to controls, while other serotypes showed a similar but non-significant trend (**Figure 1E**).

Aspartate aminotransferase (AST) and alanine transaminase (ALT) are liver enzymes commonly used as indicators of liver pathology. In male BALB/c mice, normal AST and ALT levels are 99.4□±□39.6 U/L and 135.2±26.53 U/L, respectively.(19) In set 1 experiments, where mice received DV2 NS1 plasmid via IP (with/without FuGENE), IM, or IV route, a slight rise in mean AST level was observed in DV2 NS1 administered mice compared to combined control groups (negative and FuGENE control) (**Figure 1F**). Overall, serum AST levels appeared slightly elevated in IM and IV groups compared to the combined control group. However, these differences were not statistically significant (**Figure 1G**).

The mean ALT level showed minimal variation in DV2 NS1 administered group relative to combined controls, though a slight increase was noted (**Figure 1H**). A marked elevation in ALT was seen in the IM and IV groups, with a statistically significant increase (p<0.01 and p<0.05 respectively) compared to the combined controls (**Figure 1I**). Elevated ALT levels were also observed in the IM and IV groups relative to mice inoculated with DV2 NS1 plasmid via IP (with/without FuGENE) (**Figure 1I**).

In set 2, mice administered with DV1–4 NS1 via IP showed significant (p<0.001) increase in overall mean AST level compared to their respective controls (**Figure 1J**). AST levels in DV1 NS1 and DV2 NS1 groups showed statistically significant rise (p<0.05 and p<0.01 respectively) compared to their respective controls (**Figure 1K**). No change in ALT levels was observed across all DV serotypes (**Figures 1L and 1M**). A small rise in ALT level was seen in DV4 NS1 group compared to the control (**Figure 1M)**.

### DV NS1 plasmid augmented serum glucose level in mice

Serum glucose level was estimated for all the DV NS1 plasmid administered mice using Sinocare Safe AQ max I Blood Glucose Meter (SIHC GmbH, Germany). The normal range of fasting blood sugar in mice is 80-100 mg/dL and after food intake it is <140 mg/dL.(20, 21) The mean glucose level was found significantly higher (p<0.01) in all DV2 NS1 plasmid inoculated mice compared to all controls combined (n=9 mice, control groups 1 plus 2) for set 1 experiment **(Figure 2A)**. Among the different groups, an increased trend in glucose level was observed in mice administered with DV2 NS1 plasmid by different routes compared to the control group 1 (n=6 mice) or FuGENE control group 2 (n=3 mice) **(Figure 2B)**. However, the differences were not significant.

**Figure 2:**
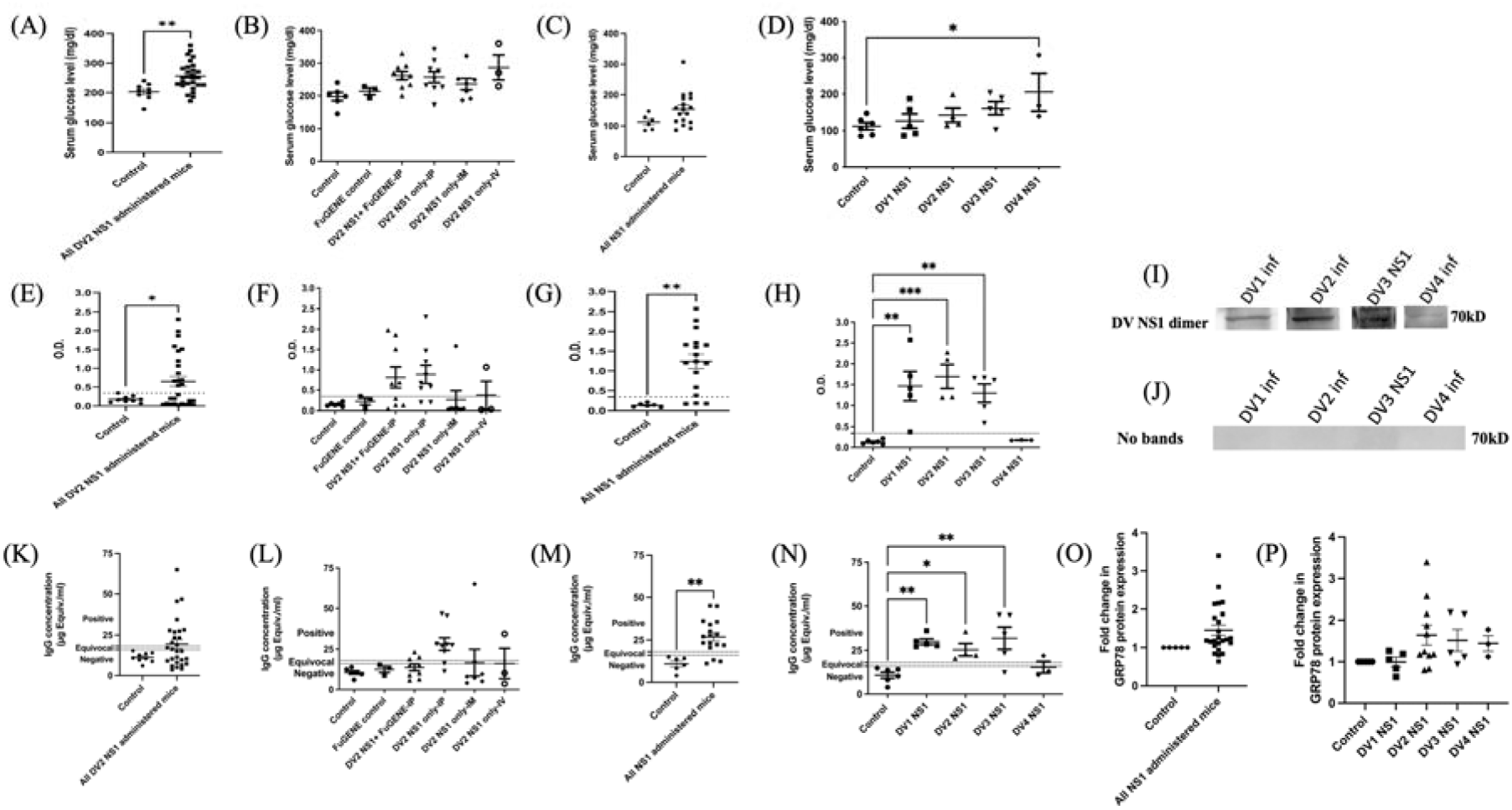
DV NS1-induced alterations in serum glucose levels, NS1-specific Ab response, and immune complex formation in mice. (A–D) Serum glucose levels in DV NS1-administered mice. (A) Mean serum glucose levels of DV2 NS1 plasmid-inoculated mice (via IP, IM, and IV routes combined) compared to control. (B) Route-specific glucose levels in DV2 NS1-inoculated mice (IP, IM, and IV). (C) Mean glucose levels in mice administered with serotype specific DV NS1 plasmids compared to control. (D) Serotype-specific glucose levels in mice administered with NS1 plasmid via IP route. **(E–J)** Detection of anti-NS1 Abs in mice serum samples. (E) Mean OD values for DV2 NS1-specific Abs in all DV2 NS1 plasmid-inoculated mice versus controls. (F) Route-specific OD values for DV2 NS1 Abs compared to control and FuGENE control groups. (G) Combined Ab responses across all four serotype-specific DV NS1 administered mice vs control group. (H) Serotype-specific DV NS1 Ab levels in plasmid-inoculated mice compared to controls. A threshold OD >0.342 indicates seropositivity, marked by horizontal dotted lines. (I) Representative WB analysis confirming the presence of NS1 Abs in sera from mice immunized with NS1 plasmid DNA. Cell lysates from DV-infected or NS1-transfected Huh7 cells were used as antigen sources. Blots were probed with serum collected from NS1-immunized mice, followed by HRP-conjugated secondary Ab. Specific immunoreactive bands corresponding to NS1 were detected, confirming Ab generation against NS1 in vivo. Lane 1: DV1-infected lysate+serum from DV1 NS1-inoculated mice (DV1A4D60); Lane 2: DV2-infected lysate+DV2 NS1 serum (DV2A1D45); Lane 3: DV3 NS1-transfected lysate+DV3 NS1 serum (DV3A2D45); Lane 4: DV4-infected lysate+DV4 NS1 serum (DV4A3D60). (**J**) Control blot included pooled sera from 3 mock-immunized mice (G1A2D14, G2A2D7 and Control mice A4D60). **(K–N)** Circulating immune complex (CIC) and aggregated IgG levels. (K) Mean aggregated IgG levels in DV2 NS1 plasmid inoculated mice (all routes combined) versus controls. (L) Route-specific CIC-C1q levels in DV2 NS1-administered mice (IP, IM, IV). (M) Aggregated IgG levels in mice administered with NS1 plasmids from all four DV serotypes compared to controls. (N) CIC-C1q levels in mice inoculated with serotype-specific NS1 plasmids via IP route. A threshold of 18 µg Eq/mL IgG indicates CIC positivity, shown by horizontal dotted lines. (**O-P**) Circulating GRP78 levels in mice. (O) Mean fold change in circulating GRP78 levels (normalized by Ponceau S staining) in NS1 administered mice serums (all serotypes combined) verses control mice serums. (P) Serotype specific fold changes in GRP78 levels compared to control. Data represented as mean±SEM. (*p<0.05; **p<0.01; ***p<0.001).

In set 2 experiment, the overall mean serum glucose level showed a higher trend in mice inoculated with serotype specific NS1 compared to the control group (n=6) **(Figure 2C)**, although it was not significant. Among the individual groups of mice receiving NS1 plasmid of four different DV serotypes intraperitoneally, the mean glucose level was significantly higher (p<0.05) in case of DV4 plasmid inoculated mice compared to the control mice **(Figure 2D)**. The mean serum glucose levels of all the groups have been cited in **Table 1**.

**Table 1:** Mean±SEM expression of DV NS1 Ab, serum glucose level and formation of immune complex in NS1 plasmid administered mice serum.

|  | Mean ± SEM<br>OD of NS1<br>Ab | NS1 Ab | Mean ± SEM<br>Serum<br>glucose level<br>(mg/dl) | Mean ± SEM<br>µg Equiv./ml<br>of aggregates<br>IgG | Formation<br>of immune<br>complex | Mean ± SEM<br>AST | Mean ± SEM<br>ALT |
| --- | --- | --- | --- | --- | --- | --- | --- |
| **Set 1 Experiment** |  |  |  |  |  |  |  |
| Control IP | 0.149 ± 0.02 | Negative | 197.5 ± 12.77 | 10.603 ± 1.10 | Negative | 663.882±67.59 | 14.914±0.652 |
| FuGENE control IP | 0.225 ± 0.08 | Negative | 213.7 ± 10.59 | 12.529 ± 1.96 | Negative | 605.431±21.25 | 13.472±0.221 |
| DV2 NS1+<br>FuGENE –IP | 0.814 ± 0.26 | Positive | 261.8 ± 13.02 | 13.780 ± 2.08 | Negative | 641.426±72.599 | 15.585±0.779 |
| DV2 NS1 only-IP | 0.889 ± 0.23 | Positive | 256.9 ± 17.00 | 28.186 ± 3.91 | Positive | 696.835±61.895 | 12.466±1.308 |
| DV2 NS1 only-IM | 0.267 ± 0.22 | Negative | 235.7 ± 17.46 | 16.655 ± 8.23 | Equivocal | 781.710±25.509 | 24.351±3.426 |
| DV2 NS1 only-IV | 0.376 ± 0.35 | Positive | 286.7 ± 38.62 | 16.166 ± 9.39 | Equivocal | 784.397±100.711 | 23.877±0.699 |
| **Set 2 Experiment** |  |  |  |  |  |  |  |
| Control | 0.132 ± 0.02 | Negative | 113 ± 10.00 | 10.886 ± 1.74 | Negative | 792.645±94.547 | 18.235±3.836 |
| DV1-NS1 IP | 1.463 ± 0.36 | Positive | 127 ± 19.65 | 29.938 ± 1.65 | Positive | 1197.12±57.718 | 26.12±5.243 |
| DV2-NS1 IP | 1.694 ± 0.29 | Positive | 143 ± 19.39 | 25.362 ± 3.51 | Positive | 1247.69±52.031 | 13.831±2.425 |
| DV3-NS1 IP | 1.294 ± 0.22 | Positive | 162 ± 17.43 | 31.922 ± 6.34 | Positive | 1099.966±74.176 | 21.253±9.797 |
| DV4-NS1 IP | 0.174 ± 0.01 | Negative | 205 ± 51.52 | 15.390 ± 3.32 | Negative | 1000.653±124.492 | 48.674±21.928 |

### Expression of DV NS1 antigen (Ag) in serum

Platelia Dengue NS1 Ag ELISA kit was used to check for the presence of NS1 Ag in the serum of representative mice from all the groups and an OD of >1 was considered positive according to manufacturer’s protocol. However, among the mice in set 1 experiments, it was found that NS1 Ag was not detectable (OD<0.5) in all the DV2 NS1 plasmid inoculated mice where NS1 was administered via IP route. On the other hand, all mice tested from the IM and IV groups were in the equivocal range (OD 0.5-1) **(Figure S1C).**

Among the mice injected with NS1 plasmid of four different DV serotypes in set 2 experiments, a few had NS1 Ag levels in the equivocal range while majority were negative **(Figure S1D)** as listed in **Table S2**.

### Expression of NS1 antibody in NS1 plasmid administered mice

Serum samples from DV NS1 plasmids-administered mice were tested individually by NS1 Abs detection ELISA. An OD of >0.342 and its corresponding fold change of >0.201 with respect to the OD of the anti-dengue virus NS1 glycoprotein Ab (DN3) was considered Ab positive. The reason behind taking OD 0.342 as cut-off was that this was the highest background OD observed in a control mouse **(Table S2)**. DN3 OD was taken as pan NS1 Ab positive control.

The mean NS1 Ab from all DV2 NS1 plasmid inoculated mice was significantly higher (p<0.05) compared to that of all controls i.e. FuGENE and non-FuGENE controls combined (n=9 mice, control groups 1 plus 2) **(Figure 2E)**, although there was no significant difference of individual groups (including FuGENE control group 2) with the control group 1 (n=6) **(Figure 2F)**. Within the IP group of set 1 experiment, which had two sub-groups (NS1 plus FuGENE and only NS1 plasmid), NS1 Abs were identified in 12 out of 18 mice combining both sub-groups. In the case of the IM and IV groups, only 1 mouse out of 7 (G5 A3 D55) and 1 mouse out of 3 (G6 A1 D7), respectively, tested positive for NS1 Abs **(Figure 2F) (Table S2)**.

The combined mean NS1 Ab level in set 2 experiment, was found significantly higher (p<0.01) in four different DV NS1 plasmid inoculated mice groups compared to the control mice (n=6) **(Figure 2G)**. Among the mice injected with different serotypes of DV NS1 plasmid, NS1 Abs were significantly increased in all the animals of DV1 (p<0.01), DV2 (p<0.001) and DV3 NS1 groups (p<0.01) compared to the control group **(Figure 2H)**. On the other hand, NS1 Ab could not be detected by NS1 Abs detection ELISA in any of the animals in the DV4 serotype group.

To further validate the presence of NS1 Abs in mice serum samples, Huh7 cells were infected with DV1 to DV4 serotypes, and Western blot was performed to detect NS1 Ag in the cell lysates. The NS1 Ag was successfully detected as specific bands for DV1, DV2, and DV4 when mice serums containing NS1 Abs were used as the primary Ab. However, in case of DV3 infection, the NS1 Ag level in the cell lysate was minimal and did not yield a detectable band. To overcome this, Huh7 cells were transfected with a plasmid encoding DV3 NS1. The resulting cell lysate, enriched in DV3 NS1 Ag, was subjected to WB, and a specific NS1 band was observed upon staining with the mice serums from DV3 NS1 plasmid inoculated mice, confirming the presence of DV3 NS1-specific Abs. Control blots were stained with pooled sera from mock-immunized mice. Representative blot images were presented in **Figure 2I and 2J**, with the corresponding full-length blots provided in **Figures S1E-I**.

### *In vivo* detection of DV NS1-plasmid DNA and RNA transcripts

A total of twelve representative liver samples were selected for DNA and RNA extraction. Six of these were from the NS1 Ab-positive IP group in the set 1 experiment—three from the NS1 plus FuGENE group and three from the only NS1 plasmid group. Additionally, one NS1 Ab-positive sample each was selected from the IM and IV groups (**Table 2**). DNA samples were used directly for PCR analysis, while RNA samples were treated with DNase to eliminate any residual DNA. Subsequently, reverse transcription PCR (RT-PCR) was carried out on the DNA-free RNA.

**Table 2:**
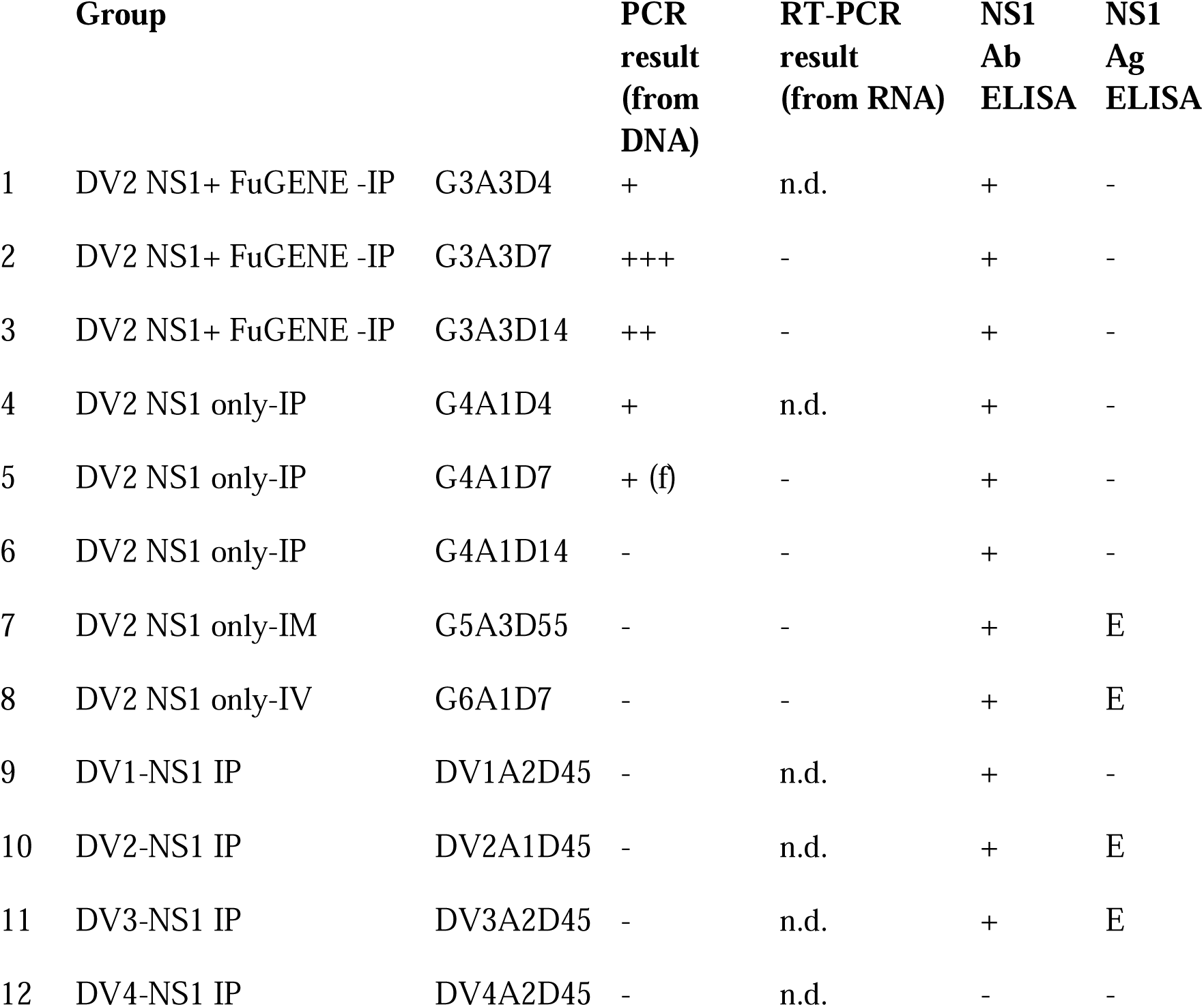
Detection of NS1 DNA in liver samples. (G=Group, A=Animal, D=Day of sacrifice, IP= Intraperitoneal, IM= Intramuscular, IV= Intravenous, E= Equivocal and n.d.=not done). Data represented as (+) for presence of DV2 NS1 plasmid DNA in NS1 administered mice liver samples and NS1 Ab in mice serums, (-) for absence of DV2 NS1 plasmid DNA/RNA in NS1 administered mice liver samples and NS1 Ag in mice serum samples, E stands for equivocal range of NS1 Ag in mice serum samples and (f) stands for presence of faint band in agarose gel for DV2 NS1 plasmid DNA in NS1 inoculated mice liver (**Figure S1I-L**). For further details, please see Table S2.

Gel electrophoresis of the PCR products revealed distinct DV2 NS1-specific bands in all three samples of the NS1 plus FuGENE (G3) group (**Table 2**). In contrast, the only NS1 plasmid group (G4) showed faint bands in two of three samples while no bands were detected in the IM (G5) or IV (G6) groups (**Figure S1J**). Similarly, no bands were observed in liver samples from DV serotype-specific NS1 plasmid-injected groups, even at 45 days post-inoculation (**Figure S1M**). Furthermore, no RT-PCR product bands were detected in any RNA samples from set 1 across all groups, indicating an absence of detectable NS1 RNA (**Figures S1K-L**).

### Formation of immune complex in DV NS1 plasmid administered mice

Serum samples from DV NS1 plasmid administered mice were individually assessed for the formation of immune complex (IC) using the CIC-C1q ELISA kit, where the IgG concentration above 18 μg Equiv./ml indicated a positive result.

No significant difference was observed in the mean IC level among all DV2 NS1 plasmid inoculated mice by different routes compared to the controls combined (n=9 mice, control groups 1 plus 2) **(Figure 2K)**. In the IP group of set 1 experiment, which was divided into two sub-groups (NS1 plus FuGENE and only NS1 plasmid groups), IC formation was detected in 3 out of 9 and 7 out of 9 mice respectively, across both sub-groups. For the IM and IV groups, only 1 mouse out of 7 (G5 A3 D55) and 1 out of 3 mice (G6 A1 D14), respectively, tested positive for CIC C1q ELISA **(Table S3)**. An increasing tendency in IgG concentration was observed in the NS1 plasmid (without FuGENE) inoculated IP group (group 4) compared to the control (group 1) **(Figure 2L)**.

In set 2 experiment, the mean CIC-IgG complex level was found significantly higher (p<0.01) in four different DV NS1 plasmid administered mice combined compared to the controls (n=6) **(Figure 2M)**. Amongst the mice which were injected with DV serotype-specific NS1 plasmid, all animals in the DV1 and DV2 groups showed the presence of IC. All but one mouse of the DV3 NS1 group of animals tested positive for CIC C1q ELISA; only the DV3 A4 D60 mouse was negative among the DV3 NS1 group. On the contrary, only one mouse showed IC formation among all the DV4 NS1 plasmid inoculated mice (DV4 A1 D45). Concentration of C1q IgG IC was significantly elevated compared to controls for serotypes 1, 2 and 3 **(Figure 2N).**

### Circulating GRP78 levels in NS1 administered mice

Representative serum samples from DV NS1 plasmid administered mice were analysed for circulating GRP78 by WB using a 1:20 dilution across all groups. Normalization was done using Ponceau S stain. While no statistically significant differences were detected between serotype-dependent individual NS1-administered groups and control mice, pooled analysis revealed a consistent trend towards elevated GRP78 level in NS1-treated animals (**Figure 2O**). Serotype-specific comparisons showed that DV2 NS1, DV3 NS1, and DV4 NS1 groups exhibited higher GRP78 levels relative to controls, whereas DV1 NS1 mice displayed levels comparable to baseline (**Figure 2P**). The WB images of GRP78 expression in serums of controls and NS1 inoculated mice were shown in **Figure S1N**.

### DV replication in pancreatic beta cells and GAPDH expression in liver cells

#### DV growth curve in mouse pancreatic beta cell line

In this study, a laboratory strain of DV2 (DV2_LS) was inoculated into Min6 cells. The infection resulted in an exponential increase in virus titres in the extracellular supernatant over time, while the intracellular levels increased steadily up to 120 hours and remained nearly constant between 120 to 168 hours post-infection (hpi). The viral titre in the extracellular supernatant (EC) increased 1,000-fold (3 log10 change), from 1.52 × 10□ ± 4.6X10^3^ copies at 48 hours to 3.07 × 10□ ± 5.7X10^6^ copies at 168 hpi. Similarly, the intracellular (IC) viral titre showed a 100-fold increase (2 log10 change), rising from 4.33 × 10□ ±5.6X10^3^ RNA copies at 48 hours to 2.74 × 10□ ±2.7X10^5^ copies at 168 hpi. The results also showed a progressive increase in cell death, reaching around 15% by 120 hpi followed by 48% at 144 hpi. Furthermore, at 168 hpi, there was a significant increase in cell death, reaching approximately 77%. This possibly led to the release of large quantity of intracellular virus particles into the extracellular environment, causing a sharp increase in the extracellular viral titres at this time-point. Despite these changes, the intracellular viral titres increased with time, suggesting that DV2_LS replication was maintained within the cells until cell death and subsequent lysis occurred, releasing the virus particles in the surrounding medium. The highest viral titres in both supernatant and inside the cells were detected at 168 hpi **(Figure 3A)**.

**Figure 3:**
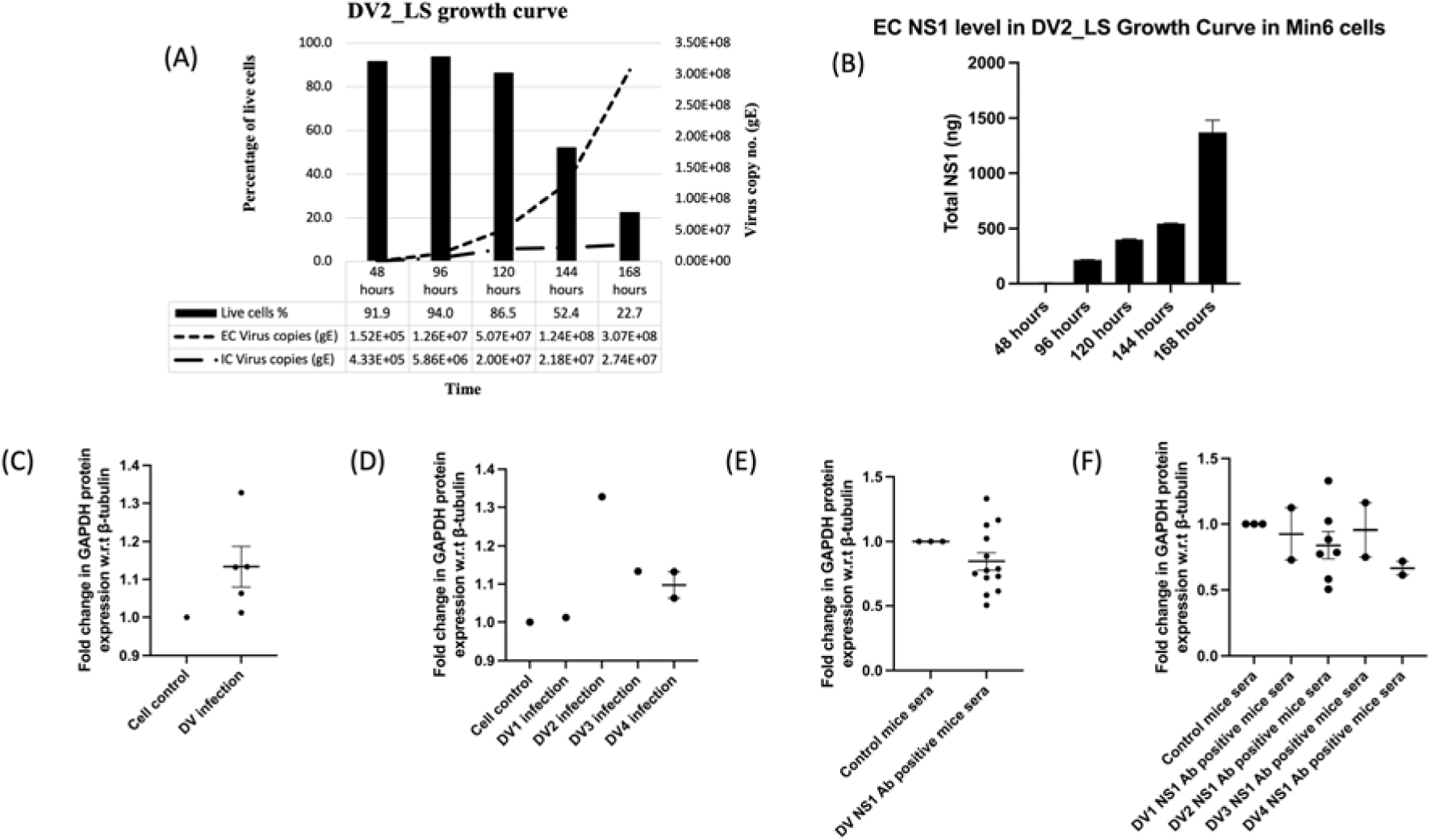
DV infection in pancreatic β-cells and serotype-dependent modulation of GAPDH expression in hepatic cells. **(A–B)** DV2 infection kinetics in pancreatic β-cell line. (A) Min6 cells were infected with DV2_LS, and samples were collected at indicated time points. Viral genome equivalents (gE) were quantified in extracellular (EC, 1□mL supernatant) and intracellular (IC, 300□µL lysate) fractions using qRT-PCR. Trypan blue exclusion was used to assess cell viability. Column graphs represent the percentage of live cells; line graphs depict viral load over time. (B) Total secreted NS1 levels (ng/mL) at different time points in the supernatant of infected Min6 cells were measured via quantitative NS1 ELISA. **(C–D)** GAPDH protein expression following DV infection. (C) Mean fold change in GAPDH expression (normalized to β-tubulin) in DV-infected Huh7 cells (all serotypes combined) versus uninfected controls at 96 hpi. (D) Serotype-specific fold changes in GAPDH expression in Huh7 cells infected with individual DV serotypes at 96 hpi. **(E–F)** Effect of NS1 Ab on GAPDH expression in hepatic cells. (E) Mean fold change in GAPDH expression in Huh7 cells treated with NS1 Ab-positive sera compared to control mouse sera at 96 hours. (F) Fold changes in GAPDH expression following treatment of Huh7 cells with NS1 Ab-positive sera from different DV serotype-specific NS1 administered mice compared to control sera at 96 hours. Data represented as mean±SEM.

The EC NS1 levels were monitored at multiple time points following DV2_LS infection in Min6 cells, revealing a progressive and sustained increase indicative of active viral replication. A substantial 33-fold rise in NS1 concentration was recorded between 48 and 96 hpi. In contrast, the increases between 96- to 120 hours and 120- to 144 hours were more modest, showing approximately 1.9-fold and 1.4-fold elevations, respectively. Notably, a second surge in NS1 secretion occurred between 144- and 168 hours, with levels increasing by 2.5-fold. These data collectively demonstrated a time-dependent escalation in NS1 secretion, reaching a peak level of 1369 ± 111.8 ng at 168 hpi, thereby providing further evidence of productive DV2_LS replication in Min6 cells **(Figure 3B)**.

### DV infection in mammalian liver cell line affected GAPDH expression

In the previous set of experiments, it has been demonstrated that DV can replicate in pancreatic β-cell line (Min6). Building on this, we aimed to investigate whether GAPDH, a key regulatory enzyme in glycolysis, was impacted during infection. To address this, Huh7 cells were infected with all four DV serotypes: DV1 and DV3, non-cytopathic clinical isolates, and DV2_LS and DV4_LS, non-cytopathic and cytopathic laboratory strains, respectively.

In DV-infected Huh7 cells, the mean GAPDH protein level across all four serotypes combined was higher than that of the uninfected controls at 96 hpi, although the difference was not statistically significant (**Figure 3C**). When analysed individually, all four serotypes exhibited a trend toward increased GAPDH protein expression compared to the control, using β-tubulin as the reference protein (**Figure S1O**). Among the serotypes, DV2 showed the maximum increase in GAPDH expression, followed by DV3, DV4, and DV1. However, none of these increases reached statistical significance (**Figure 3D**).

To confirm that all four DV serotypes successfully replicated in Huh7 cells, IC NS1 level was assessed by Western blot (**Figure S1O**).

### DV NS1 Ab treated liver cells showed reduction in GAPDH protein expression

Huh7 cells were treated with serum samples derived from DV NS1 Ab-positive mice and incubated for 96 hours. A noticeable, though statistically non-significant, reduction in mean GAPDH protein level was observed in Huh7 cells treated with serum from mice injected with NS1-expressing plasmids of various DV serotypes, compared to cells treated with control mice serum samples. For the WB analysis, β-tubulin was kept as the house-keeping reference control (**Figure S1P**). Notably, 69% of the NS1 Ab-positive mice serums, when incubated with Huh7 cells for 96 hours resulted in decrease in GAPDH expression (**Figure 3E**).

In case of set 2 experiments, two representative serum samples for each serotype of DV NS1 plasmid administered groups were selected for analysis. Additionally, from set 1 experiment, two serum samples from the NS1 plus FuGENE group (G3) and three from the NS1-only group (G4) were randomly chosen. These five samples of G3 & G4 were plotted with the DV2 NS1 group from set 2 for comparative analysis.

When the mean GAPDH protein expression was evaluated individually across the four serotypes of DV NS1 Ab-positive serum-treated Huh7 cells, a downward trend was observed relative to the control, although none of the differences reached statistical significance. Among the NS1 Ab-treated groups, the greatest reduction in GAPDH expression was observed in cells exposed to DV4 NS1-positive serums, followed by DV2, DV1, and DV3 (**Figure 3F**).

### Histopathology revealed several cytological anomalies

Histopathological slides were prepared for liver, spleen, lungs, heart, and brain from the DV2 NS1 IP group (set 1). For DV2 NS1 IM and IV groups (set 1) and serotype-specific DV NS1 groups (set 2), only liver and spleen sections were analyzed.

In the DV2 NS1 IP group of set 1, liver sections showed mononuclear infiltration (green arrow) at 4- and 7-days p.i. Ballooning of hepatocytes (yellow arrows) began at day 7 and peaked at day 14, along with portal congestion in FuGENE□+□NS1 administered mice, indicating fibrosis (**Figures 4A and S2A**).

**Figure 4:**
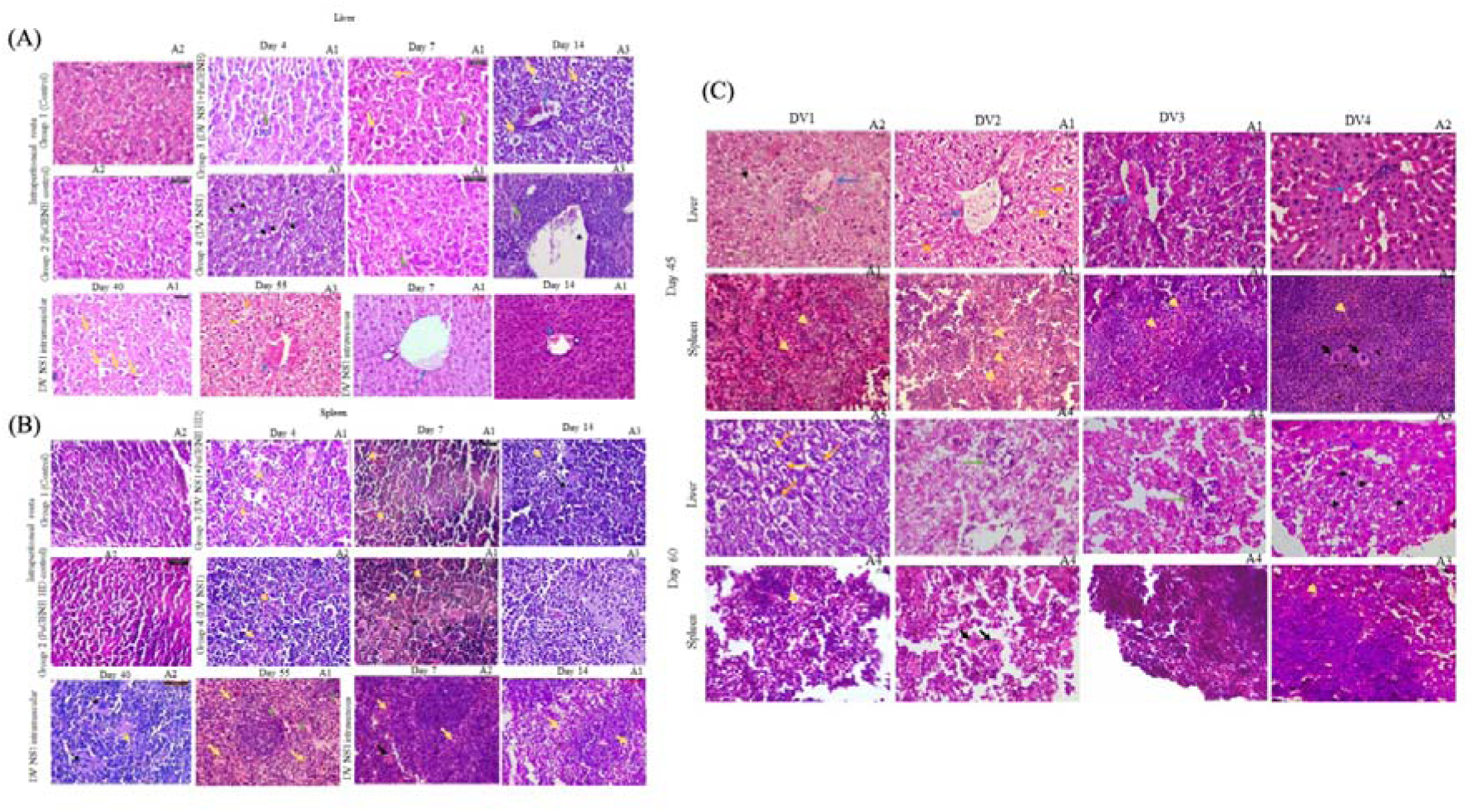
Histopathological analysis of liver and spleen in DV NS1 plasmid administered mice. **(A)** Representative hematoxylin and eosin (H&E) stained liver sections from DV2 NS1 plasmid-injected mice via IP, IM, and IV routes. Control and FuGENE control groups exhibited normal hepatic architecture. DV2 NS1 treated groups showed central vein congestion (blue arrow), mononuclear infiltration (green arrow), ballooning degeneration of hepatocytes (yellow arrow), and fat globule accumulation (black arrow). **(B)** H&E stained spleen sections from the same groups revealed normal lymphocyte distribution in control and FuGENE control mice. In DV2 NS1 treated mice, severe lymphocyte depletion in the white pulp (yellow arrow), apoptotic bodies (green arrow), and macrocytic megakaryocytes (black arrow) were observed. A foamy appearance was noted in spleens collected at day 14 post-IP administration. **(C)** Serotype-dependent pathology following DV NS1 administration. Liver sections from mice administered with serotype-specific DV NS1 plasmids (DV1–DV4) via IP route and sacrificed at day 45 showed central vein congestion (blue arrow) and hepatocyte ballooning (yellow arrow). Spleen sections at day 45 showed lymphocyte depletion across all serotypes. Both liver and spleen sections exhibited extensive tissue damage by day 60 in all the serotypes with liver sections showing hepatocyte ballooning (yellow arrow), mononuclear infiltration (green arrow), and mild steatosis (black arrow). All images were captured at 40X magnification.

Mice receiving DV2 NS1 via IM route exhibited hepatocyte degeneration and ballooning at day 40 p.i., and severe central vein congestion (blue arrow) at day 55, suggesting progressive liver damage. In the IV group, mild hepatic portal vein congestion at day 7 p.i. worsened by day 14 (blue arrow).

Spleen sections from DV2 NS1 IP groups showed lymphocyte depletion in the white pulp (yellow arrow), with circular macrophages and apoptotic bodies visible at days 4 and 7, and peak damage at day 14 (black arrow). Foamy lymphocytes appeared on day 14 in the DV2 NS1-only group. IM group spleens showed similar lymphocyte depletion, apoptotic bodies (green arrow), and megakaryocytes (black arrow). IV group spleens also exhibited lymphocyte depletion at days 7 and 14. No other significant histopathologic changes were noted (**Figures 4B and S2B**).

In set 2, liver sections of all DV NS1 groups showed central vein congestion (blue arrow) and mononuclear infiltration (green arrow). By day 60, liver tissue architecture was severely disrupted, with no clear hepatocyte demarcation. Steatosis was evident in the DV4 NS1 group at day 60 (black arrow). Livers collected at day 60 were harder and more brittle than those at day 45 (**Figures 4C and S2G**).

Spleens from all serotype-specific DV NS1 groups showed splenocyte depletion (yellow arrow). Multiple megakaryocytes (black arrow) were observed particularly in DV2 NS1 (day 60) and DV4 NS1 (day 45) groups. Morphologically, spleens were enlarged, indicating splenomegaly (**Figures 4C and S2H**).

Lung section of DV2 NS1 plasmid administered mice via IP route from set 1 experiment indicates atelectasis (black circle) and alveolar fusion (black star) at 7 days p.i. compared to sham control and vehicle (FuGENE) control groups. Similar damage was seen in the IM group lung sections in the form of atelectasis (black circle) at day 40 **(Figures 5A and S2C)**.

**Figure 5:**
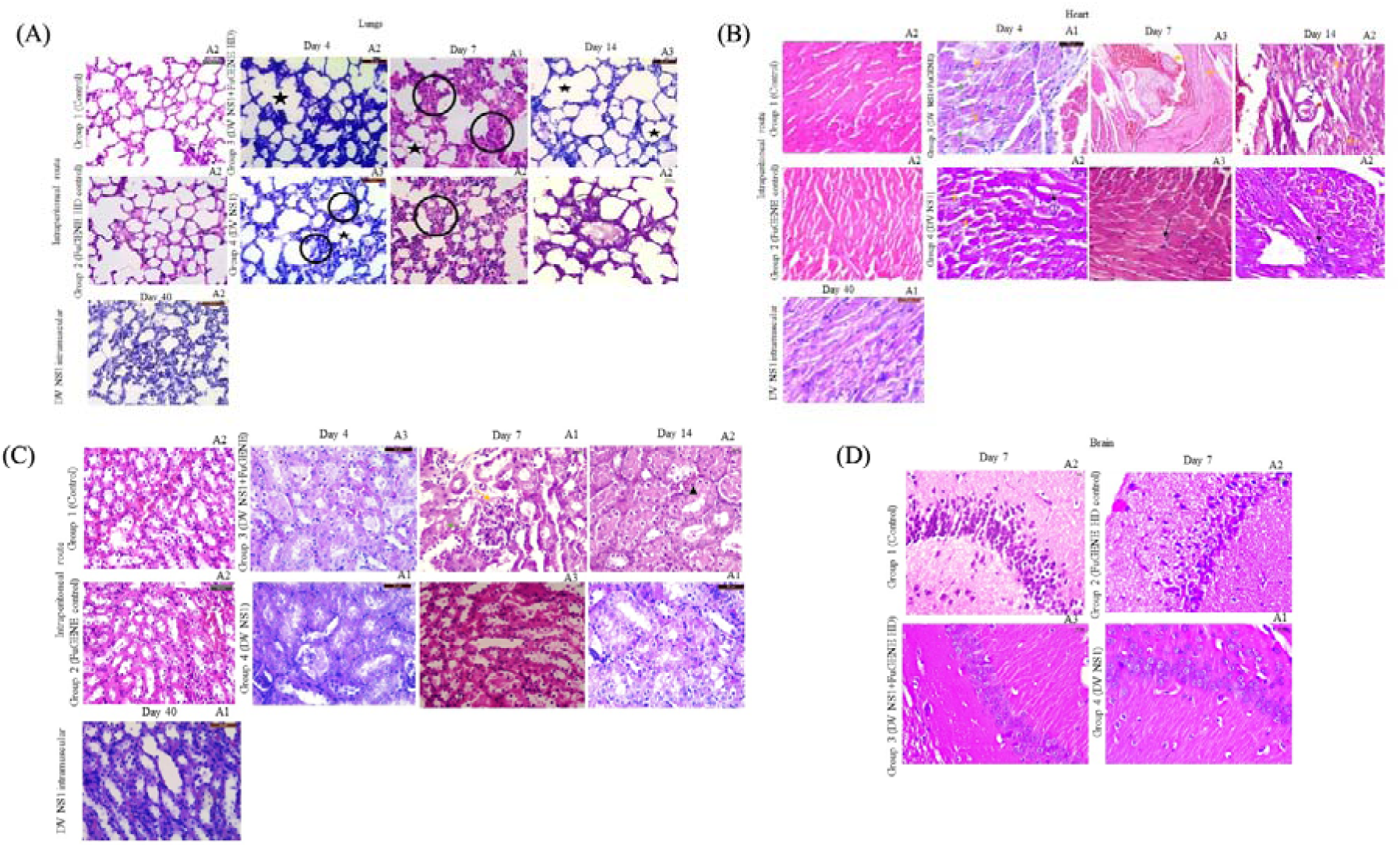
Histopathological alterations in non-hepatic tissues of DV2 NS1 plasmid administered mice. **(A)** H&E stained lung sections from DV2 NS1 plasmid inoculated mice (IP and IM routes) showed structural damage, including alveolar atelectasis (black circle) and alveolar fusion (star). Control and FuGENE control groups demonstrated normal alveolar architecture. **(B)** Cardiac muscle section of control, FuGENE control and NS1 IM group showed normal histology. Interstitial edema (yellow arrow), perivascular fibrosis (green arrow) and cardiac muscle mononuclear infiltration (black arrow) was observed in DV2 NS1 administered groups via different routes. **(C)** Kidney sections from FuGENE-mediated intraperitoneal DV2 NS1-injected mice exhibited mild renal damage characterized by hyaline cast deposition (black arrow), atopic tubular degeneration (yellow arrow), and interstitial edema (green arrow). **(D)** Brain section of DV2 NS1 IP-injected mice showed diffuse neuronal loss in the CA1 region of the hippocampus at day 7 post-inoculation. Control and FuGENE HD control groups presented densely packed intact pyramidal neurons. All images were captured at 40X magnification.

Heart muscle histopathology from set 1 experimental mice indicated cardiomyocyte edema (yellow arrow) and mononuclear infiltration indicating inflammation (black arrow) in both the DV2 NS1 IP subgroups. Maximum cardiac degeneration was observed after day 14 in FuGENE mediated DV2-NS1 plasmid administered group with vascular leakage, arteriolar detachment from peripheral tissues (red arrow). Perivascular fibrosis was noted at day 4 in case of DV2 NS1 plasmid administration. No visible sign of damage has been observed in cardiac tissue via IM route **(Figures 5B and S2D)**.

Kidney of set 1 experiment mice from FuGENE mediated DV2-NS1 plasmid administered group via IP route showed hyaline cast inside the lumina of renal tubules (black arrow) and atopic degeneration of renal tubules (yellow arrow) along with edema (green arrow) at 7 days p.i.. No significant renal tissue damage was observed in the DV2 NS1 IM group **(Figures 5C and S2E)**.

Brain section of mice from set 1 experiment injected with DV2-NS1 plasmid via IP route, was analysed on day 7 and the hippocampus region was observed. CA1 region of DV2 NS1 group, with and without FuGENE, manifested diffuse neurons compared to sham control and FuGENE control groups which showed intact neurons **(Figures 5D and S2F)**.

## Discussion

DV continues to be the most critical and neglected tropical viral disease globally. Among its proteins, NS1 stands out as the most abundantly secreted Ag with diverse roles in virus replication, immune modulation, and pathogenesis. NS1 has been linked to endothelial dysfunction, immune activation, and tissue injury, underscoring its importance in host–virus interactions.(22) In this study, we investigated the effects of NS1 administration in a murine model.

To identify an optimal delivery route, BALB/c mice were injected with DV2 NS1 plasmid via IP, IM, or IV route. IP delivery was prioritized due to its unique pharmacokinetics-bypassing gastrointestinal metabolism while still involving hepatic first-pass processing.(23) Furthermore, lipophilic transfection agents such as FuGENE, had been known to improve IP-mediated systemic uptake.(24) However, comparative evaluation of IP, IM and IV routes revealed that IP administration (without FuGENE) generated consistent Ab responses, thereby serving as the preferred delivery mode for subsequent serotype-specific investigations. A prime–boost strategy using 100 µg/mouse of DV NS1 plasmid via the IP route, consistent with published protocols, effectively induced humoral responses.(25–28) The prime–boost regimen (days 1, 18, 33) with 100 µg elicited stronger Ab and immune responses than the low-dose, three-day consecutive schedule (20 µg on days 1–3).(25)

Previous reports on NS1 plasmid-based immunization have largely emphasized on Ab titres, often without detecting circulating NS1 antigen post-inoculation.(18,25,27) Our results aligned with these observations. The transient presence of episomal plasmids likely restricted antigen expression to a narrow temporal window, precluding sustained antigenemia but allowing sufficient antigen presentation to elicit adaptive immune responses.

Distinct NS1 PCR bands were detected in liver DNA after IP delivery, particularly with FuGENE, whereas for IM and IV routes, NS1 plasmid was not detectable by PCR in liver DNA. DV2-NS1 expression (set 1 experiment) was associated with slightly elevated AST and ALT levels. Set 2 experiments revealed a significant AST increase and a trend towards higher liver weight, while ALT remained comparable to controls. Additionally, spleen weight was significantly increased in serotype-specific NS1–administered mice.

Alongside its immunological implications, DV infection significantly affects host metabolism. Pre-existing hyperglycaemia has been linked to worsened disease outcomes and enhanced virus replication.(29–31) Interestingly, there are few case reports and cohort studies that highlighted that DV infection was associated with alterations in insulin sensitivity and glycemic control, often accompanied by pancreatitis.(32–37) Mechanistically, DV3 infection had been shown to downregulate insulin receptor substrate-1 (IRS-1) in hepatocytes via TNFα signalling, promoting insulin resistance.(38) These findings raised important questions about the immune-metabolic interplay in dengue pathogenesis.

SARS-CoV-2 can infect pancreatic tissues via ACE2, leading to β-cell dysfunction and new-onset diabetes.(39–41) Such infections have underscored the pancreas as a viral target organ with profound metabolic consequences.(42–44) Whether DV can similarly perturb β-cell function via direct infection or through immune-mediated mechanisms remained an open question. This has been addressed in the present study.

A potential mechanistic link may lie in DV-induced metabolic reprogramming. NS1 has been shown to enhance glyceraldehyde-3-phosphate dehydrogenase (GAPDH) activity during early infection to support glycolytic flux, while NS3 antagonizes this process at later stages.(45–47) We have also observed in the present study that DV infection resulted in increased GAPDH levels at 96 hpi in agreement with previous observations.

Now, GAPDH dysfunction/downregulation is a hallmark of diabetic complications and may contribute to long-term metabolic impairments.(48, 49) Notably, NS1-specific Abs can persist in circulation for extended periods(50) and our current study revealed that these NS1 Abs could potentially suppress GAPDH activity over time, suggesting a probable link between NS1 immunity and post-infection dysglycemia.

Complement activation is another cornerstone of DV immunopathogenesis.(51) While classical (C1q-IgG) and lectin pathways facilitate viral clearance via opsonization and phagocytosis, uncontrolled activation contributes to systemic inflammation and cytokine storms.(52–54) Circulatory Immune complex (IC) deposition, a central feature of type III hypersensitivity, has been associated with DV-related glomerulonephritis and other organ-specific damage.(55–58) In secondary dengue infection, rapid clearance of NS1 due to rising anti-NS1 IgG suggested early immune complex formation, which might contribute to severe disease pathogenesis.(59) Consistent with these findings, we observed IC formation in around 80% of mice administered with DV2 NS1 (set 1 experiments) or NS1 plasmid from four distinct DV serotypes (set 2 experiments).

An interesting observation was that 49% of the NS1 administered mice tested positive for both NS1 Abs and IC formation, while 13% were positive for NS1 Abs only but negative for ICs. In contrast, 11% of the NS1 administered mice were positive for ICs but negative for NS1 Abs, and the remaining 27% were negative for both. These findings suggested that NS1 Abs tend to form ICs.

Musculoskeletal complications, such as arthralgia and arthritis, have also been reported post-dengue.(60) Cases of dengue-associated sacroiliitis may be attributable to IC deposition in joint tissues, mirroring mechanisms observed in HCV-, HBV-, and parvovirus-associated arthritis.(61, 62) Though definitive experimental validation remains lacking, such manifestations suggest a systemic extension of DV-induced immunopathology.

Again, GRP78, a central endoplasmic reticulum (ER)-resident chaperone, is upregulated during DV infection as viral protein accumulation induces ER stress and activates the unfolded protein response (UPR).(63) Although primarily ER-resident, GRP78 can translocate to the cell surface or be secreted into circulation under stress, where it facilitates DV entry and supports NS1 folding and secretion.(64–66) Notably, NS1 and GRP78 physically interact, and reductions in GRP78 impair NS1 biogenesis and secretion, underscoring their interdependence.(67)

In our study, NS1 administration alone was sufficient to elevate circulating GRP78 in mice, indicating that NS1 can trigger ER stress responses. This effect may be amplified by NS1-directed Abs, which could exacerbate cellular stress and promote GRP78 release, potentially fostering GRP78 autoantibodies (auto-Abs) generation and subsequent formation of GRP78-immune complexes. Beyond dengue, GRP78 auto-Abs had been implicated in other immune-mediated diseases, including rheumatoid arthritis and type 1 diabetes mellitus (T1DM),(66, 68, 69) while GRP78 upregulation was also associated with type 2 diabetes (T2D).(70) As dysregulated UPR and chronic ER stress contribute to both autoimmunity and metabolic disease, NS1-driven GRP78 release and immune complex formation may represent a mechanistic link between dengue and diabetes-related pathologies.(71)

Anti-NS1 Abs have been shown to cross-react with platelets and endothelial cells, raising the possibility of β-cell targeting and autoimmune sequelae (type II hypersensitivity).(72–75) This hypothesis aligns with literature linking enteroviral infections to T1DM via β-cell destruction.(76, 77) Given DV’s capacity to induce auto-Abs and the autoimmune basis of T1DM, dengue infection may trigger NS1 Ab-mediated β-cell destruction (type II hypersensitivity), impairing insulin production and promoting hyperglycemia or T1DM onset. However, direct evidence is still lacking.

Furthermore, circulatory IC formation involving NS1-specific IgG may drive type III hypersensitivity responses within the pancreas, potentially contributing to latent autoimmune diabetes in adults (LADA).(78)

In this respect, the present study showed that DV could indeed, infect, replicate in (and potentially damage) the pancreatic beta cell line (Min6). The NS1 plasmid administration in mice model revealed that DV NS1 Abs could be robustly generated by this approach. Such Abs had been already proven to have autoimmune characteristics to cause direct tissue damage (evident from previous reports). Finally, our study showed that ICs were increasingly formed in the serum of NS1 plasmid administered mice which could also lodge into the tissues and damage them.

Our histopathological findings support the concept of NS1-induced tissue injury in the liver, spleen, and cardiopulmonary systems. The liver, being central to complement protein synthesis and immune surveillance, displayed mononuclear infiltration and hepatocellular degeneration, consistent with prior reports of DV-associated liver dysfunction.(79–81) Passive and active NS1 immunization models have previously demonstrated hepatic injury even in the absence of antigenemia.(82) Similarly, spleens from NS1-inoculated mice showed lymphoid depletion and inflammation, aligning with observations in patient tissues.(83, 84)

Cardiopulmonary pathologies including myocardial edema and alveolar fibrosis are increasingly recognized in dengue and were reproduced in our NS1 exposure model.(85–87) These manifestations likely arise from sustained cytokine signalling and immune activation, possibly aggravated by persistent NS1-specific IgG.

Notably, renal tissues exhibited minimal pathological changes, corroborating prior studies indicating limited kidney involvement in DV infection.(88) However, systemic effects of long-term NS1 Ab responses should not be underestimated. For example, passive transfer of IgG from long COVID patients has been shown to induce neurological pathology in murine models, and similar mechanisms might be at play in dengue.(89, 90)

The detection of hippocampal CA1 neuronal degeneration following NS1 exposure adds a novel dimension to our understanding of dengue-associated neurotropism. Neurological complications, including cognitive impairment and encephalitis, have been reported in DV patients.(91, 92) Given that CA1 degeneration is a hallmark of hippocampal sclerosis of aging (HS-A), our findings suggest that chronic NS1 Ab responses may have neurodegenerative consequences, emphasizing the need for long-term neurological surveillance in dengue survivors.(93)

DV NS1 is also a compelling antiviral target.(15) Compounds such as Ivermectin, lowered circulating NS1 levels by inhibiting importin-mediated nuclear transport of transcription factors needed for chaperones (like GRP78) that support NS1 folding and secretion.(67, 94, 95) Domperidone, on the other hand, directly interacts with viral proteins including NS1 to inhibit virus production and NS1 release, demonstrating the feasibility of this approach.(96) Our animal model is particularly suited for evaluating such candidates, as the NS1-driven pathogenic effects observed in our system would be mitigated by effective inhibitors, making it a valuable platform for preclinical antiviral screening.

In conclusion, our findings highlight the multifaceted role of NS1-specific Abs in DV pathogenesis. NS1 expression in vivo induced robust serotype-specific humoral responses accompanied by IC formation and metabolic alterations (**(Figure 6)**. Notably, NS1 Abs persist for extended durations, and the potential for Ab-mediated tissue injury via both type II and type III hypersensitivity–like reactions warrants caution. These observations provide important insights into the interplay between viral proteins, host immunity, and metabolic regulation, emphasizing the need to further dissect NS1-driven mechanisms in disease progression.

**Figure 6:**
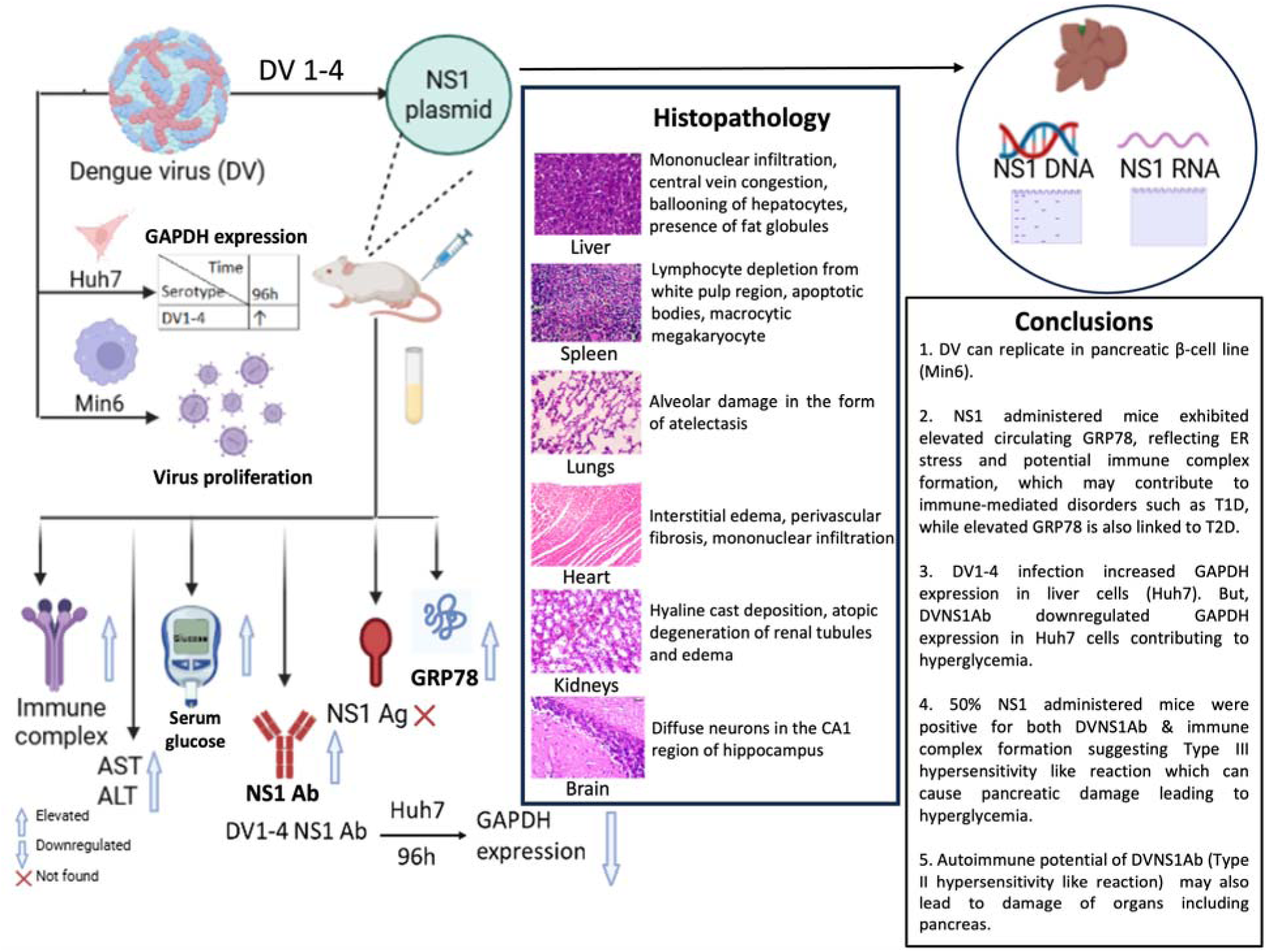
Schematic of NS1 Ab driven immune-metabolic dysfunction in DV NS1 plasmid administered mice. NS1 plasmid DNA administration in mice induced robust NS1-specific Ab generation, despite undetectable serum NS1 antigen. IP delivery caused the strongest pathology, including elevated circulating GRP78 levels, hepatic injury, splenocyte depletion, hyperglycemia and IC formation. DV replicated in pancreatic β-cells, while NS1 Ab treated liver cells altered GAPDH expression, linking NS1 Ab responses to immune-metabolic dysregulations.

## Material and methods

### DV NS1 plasmid preparation

pcDNA3.1(+) based plasmid was constructed encoding the NS1 sequence from DV1-4. The viral RNA was extracted using High Pure Viral Nucleic acid extraction kit (Roche, Germany). The RNA was used as template for the synthesis of cDNA using target gene specific reverse primer and performed by reverse transcription using Superscript III (Invitrogen, MA, USA). The segment containing the NS1 sequence was amplified by PCR using the serotype specific primers listed in **Table S1**. The PCR product was confirmed by 1% agarose gel electrophoresis and purified using PCR purification kit (Qiagen, Germany). The gene corresponding to signal sequence of 23-29 amino acids upstream of NS1 protein was included for amplifying NS1 gene necessary for cloning purpose. BamH1 and HindIII for DV1, NheI and XhoI for DV2, BamH1 and XhoI for DV3 and HindIII and EcoR1 for DENV4 restriction sites were incorporated in the forward and the reverse primers respectively. Respective restriction enzymes for each serotype were used for carrying out restriction enzyme (NEB) digestion for the PCR products and the 5.4 kb mammalian expression vector pcDNA3.1(+). Ligations were done using T4 DNA ligase (Promega, CA, USA), followed by transformation of XL1-Blue cells. Plasmids were isolated from transformed *Escherichia coli* and purified by Qiagen Plasmid Mega Kit (Qiagen, Germany), following manufacturer’s instruction. Plasmid DNAs were eluted in nuclease free water and stored at -20°C until use. DNA concentrations were determined using optical density measurements at 260nm and the integrity of the plasmids were checked by agarose gel electrophoresis. The DV1-, DV2- and DV3-NS1 recombinant plasmids were confirmed by sequencing and submitted to GenBank (Accession Numbers: MT072226, MT072227 and MT072228 respectively). DV4_LS, a laboratory strain (ATCC VR-1490), had its complete genome sequence available in GenBank under the H241 strain (Accession Number: AY947539). Accordingly, the alignment of the recombinant plasmid has been presented in **Figure S3.**

### Cell Lines

Min6, Huh7 and C6/36 cell lines were obtained from National Centre for Cell Science (NCCS), Pune, India. Min6 and Huh7 cells were cultured in DMEM (D5796, Sigma-Aldrich, MA, USA) supplemented with 10% foetal bovine serum (FBS) (Gibco, MA, USA) and Pen-Strep and L-Glutamine mix (Sigma-Aldrich, MA, USA) at 37^0^C with 5% CO_2_. C6/36 cells were cultured in MEM media (Sigma-Aldrich, MA, USA), supplemented with 10% FBS and antibiotics at 28^0^C with 5% CO_2_.

### Virus

DV2_LS (ATCC VR-1810) and DV4_LS (ATCC VR-1490) were obtained from American Type Culture Collection (ATCC, USA). DV1 and DV3, two clinical isolates were cultured as previously described.(97) For virus stock preparation, DV1, DV2_LS and DV3 were inoculated in monolayer of C6/36 cells and incubated for five days. These three viruses did not show any plaque formation, like other low-passage clinical isolates reported previously.(98) DV4_LS stock was prepared in Huh7 cells and showed cytopathic effects (CPE) as mentioned by ATCC.

### Experimental animals

Male BALB/c mice (n=63; 4-6 weeks old) weighing 20-30g were maintained in individually ventilated cages under standard conditions of temperature (23±3°C) and relative humidity (55±5%) with 12hrs light/dark cycle. 37 mice were employed for DV2-NS1 route dependent study in set 1 experiment and 26 mice were utilised in DV NS1 serotype dependent study in set 2 experiment. Animals were fed standard laboratory chow diet and water *ad libitum* throughout the study period. All the animals were acclimatized for at least 14 days before the initiation of the experiment. The animals were maintained under proper conditions till the termination of the experiment and sacrificed at the end as per study plan to evaluate various parameters.

### Ethics statement

All experimental protocols were performed in accordance with the Committee for the Control and Supervision of Experiments on Animals (CCSEA) guidelines. The study protocol was designed based on literature review in the same area and it was approved by Institutional Animal Ethics Committee (IAEC, NIPER, Kolkata).(27) The experiments on animals comply to the ARRIVE guidelines.

### Set 1 experiments: Expression of DV2-NS1 plasmid in mice via different routes of administration

The NS1 gene from DV2 strain was cloned into pcDNA3.1(+) vector. A total of 37 mice were subjected to immunization using the NS1 plasmid DNA through different administration routes **(Figure S4A)**.

Specifically, 27 out of the 37 mice were inoculated via the IP route (20µg/mouse) for 3 consecutive days, of which negative control and FuGENE (WI, USA) vehicle control groups comprised 6 and 3 mice respectively; 9 mice received DV2-NS1 plasmid with FuGENE and rest of the 9 mice were injected only DV2 NS1 plasmid without FuGENE.

7 mice received immunization through IM route (100µg) according to Mellado-Sanchez *et al* (2010) and Costa *et al* (2007), and 3 mice were administered the dose via IV (50µg) for three consecutive days.(18, 26, 27) For the IM group, booster doses were administered on day 15 and day 30 following the first dose.

For the IP group, 9 mice were sacrificed on days 4, 7, and 14 respectively following the initial dosing, while 2 mice from the IV group were sacrificed on day 7 and the remaining 1 on day 14 after their initial dosing. For the IM group, 3 and 4 mice being sacrificed respectively on days 40 and 55 and the body weight was evaluated **(Figure S4A)**. A section of liver from all the animals were stored in RNA later for DNA and RNA extraction.

### Set 2 experiments: Expression of different dengue serotype specific NS1 plasmid in mice

Four to six-week-old BALB/c mice were subjected to IP inoculation with 100μg of NS1 plasmids of different DV serotypes diluted in 100 μl of 1x PBS. A total of 26 mice were included in the study with each serotype group comprising 5 animals and 6 animals as negative control. Each group received three doses of the respective DV NS1 plasmid or control vectors, administered two weeks apart on the 1st, 18th, and 33rd days **(Figure S4B)**. The body weights of the animals were monitored and recorded weekly throughout the experiment. 2 animals from each treated group were sacrificed on day 45 and the remaining on day 60. Serum was isolated from the blood and organs were also collected for further analysis.

### AST and ALT activity assay

AST is a pyridoxal phosphate dependent enzyme present in cytosolic and mitochondrial isoenzymes and has broader tissue distribution in the liver, cardiac muscle, skeletal muscle, kidneys, brain, pancreas, lungs, leucocytes and RBC.(99) It catalyses the conversion of aspartate and α-ketoglutarate into oxaloacetate and glutamate in gluconeogenesis. Hepatocellular injury triggers the release of AST into circulation thus AST level in blood acts as a marker for liver function.

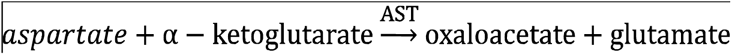

ALT is specific to hepatocytes and is an indicator of hepatocellular injury. Serum SGOT and SGPT levels were measured in DV NS1 infected mice using AST activity assay kit and ALT activity assay kit (Sigma-Aldrich, MA, USA) following the manufacturer’s instructions, respectively.

### Serum glucose estimation test

Serum glucose levels were estimated from all mice samples using the Sinocare Safe AQ max I Blood Glucose Meter (SIHC GmbH, Germany) following the manufacture’s instruction. 8.0µl of serum sample was added to the test area and the test strip was inserted in the glucometer.

### DV growth curve in Min6 cells

10^5^ Min6 cells were seeded in each well of 24-well plates. On confluency, the cells were washed with 1XPBS. The inoculum was added over the monolayer of cells at 10 MOI and incubated for 2 hours at 37°C with intermittent shaking at every 15 minutes. After 2 hours of incubation, the inoculum was removed, and the cells were washed thrice with 500 µl of 1XPBS to remove all uninternalized viruses. Then, 1 ml of DMEM supplemented with 2% FBS was added to each of the wells and the plates were incubated at 37°C for respective time-points (48-, 96-, 120-, 144- & 168-hpi). At each time-point, supernatant was collected from the wells and stored at -80°C for RNA extraction. In one of the duplicate wells, cells were trypsinized and then trypan blue staining was done to check cell viability, whereas 300 µl of fresh DMEM1 was added to the other well. RNA was extracted from 200µl of supernatant and Min6 cells using High Pure Viral Nucleic Acid Kit (Roche, Germany) and qRT-PCR was done according to our previously described protocol.(100)

### DV infection in Huh7 cells

6 × 10^5^ Huh7 cells were seeded into each well of a 6-well plate. On confluency the cells were washed with 1X PBS and infected with DV at 10 MOI for DV1, DV2_LS and DV3, and 0.04 for DV4_LS. The viral inoculum was applied to the cell monolayer and incubated at 37°C for 2 hours, with gentle shaking every 15 minutes. Following the incubation, the inoculum was removed, and the cells were washed twice with 1 ml of 1X PBS to eliminate any uninternalized virus particles. Subsequently, 3 ml of DMEM supplemented with 2% FBS was added to each well, and the plates were incubated at 37°C for 96 hours.

### Treatment of Huh7 cells with NS1 Ab positive mice serum samples

Huh7 cells were seeded at a density of 2 × 10□ cells per well in 12-well tissue culture plates. Upon reaching confluency, cells were gently washed with 500 µL of 1X PBS to remove residual media. Subsequently, 2 mL of a treatment mixture containing diluted NS1 Ab-positive mice serums (1:50 dilution) in DMEM supplemented with 2% FBS was added to each well. Cells were incubated at 37°C in a humidified atmosphere of 5% CO□ for 96 hours. Following the treatment period, cells were harvested for downstream analyses.

### DV3 NS1 transfection

2 × 10□ Huh7 cells were seeded per well in 12-well tissue culture plate. 1.0 μg plasmid DNA was transfected in each well using FuGENE HD (Promega), following manufacturer’s instructions and previously published protocol.(10)

### Western blot

Huh7 cells infected with DV or treated with NS1 Ab-containing mice serums were lysed directly on the culture plate using 400□µL and 200□µL of ice-cold lysis buffer, respectively, supplemented with a protease inhibitor cocktail (Pierce™ ThermoFischer Scientific, USA). The lysis buffer was applied to adherent cells and incubated on ice for 10 minutes. Cell lysates were then collected and centrifuged at 13,000□rpm for 15 minutes at 4□°C. The resulting supernatants were collected and subjected to protein quantification using the BCA assay (Pierce™ BCA Protein assay kit, ThermoFischer Scientific, USA). For WB with serum samples, 1:20 dilutions were made with 1X PBS and total protein concentration was determined by BCA assay. Equal amounts of protein (30□µg per sample) were prepared for electrophoresis either by heating at 95°C for 10 minutes in Laemmli buffer containing β- mercaptoethanol (β-ME) or left unheated for use with NuPAGE sample buffer. Proteins were resolved on 10% SDS–polyacrylamide gels at a constant voltage of 90□V for 2 hours, followed by transfer onto nitrocellulose membranes using the Trans-Blot Turbo Transfer System (Bio-Rad, CA, USA). Normalization for WB with mice serum was done using Ponceau S stain (GCC Biotech, India) to minimize any discrepancies in protein amount. Membranes were blocked in TBST containing 5% skimmed milk for 2 hours at room temperature and subsequently incubated overnight at 4□°C with the following primary Abs: anti-GAPDH (CST, #8884S, 1:1000), anti-GRP78 (Abcam, #ab21685, 1:1000), anti-NS1 Abs (mouse serums, diluted 1:10 or 1:100 or Abcam, #ab41616, 1:25), and anti-β-tubulin (CST, #2128S, 1:1000) as a loading control. After washing, membranes were incubated with appropriate HRP-conjugated secondary Ab for 1 hour at 4°C: anti-mouse IgG (Novus, NB7539, 1:10,000 or 1:6000) and anti-rabbit IgG (Abcam, ab97051, 1:12000 or 1:6000). Protein bands were visualized using ECL substrate (Bio-Rad) and imaged with the ChemiDoc Imaging System (Bio-Rad, CA, USA). Densitometric quantification of band intensities was performed using ImageLab software (Bio-Rad, CA, USA).

### NS1 Ag ELISA

For the detection of DV NS1 Ag in mice serum, ELISA was done as per protocol of Platelia Dengue NS1 ELISA kit (Bio-Rad, CA, USA). 50 µl of neat serums were used for ELISA and the reading was taken at 450 nm in iMark plate reader (BioRad, CA, USA).

### NS1 Ab ELISA

Detection of DV NS1 Ab was performed by ELISA using Human Anti-Dengue Virus IgG ELISA kit according to the manufacturer’s protocol with few modifications (R&D Systems, Cat# DENG00, MN, USA). Recombinant NS1 Ag of DV serotype 1, 2, 3 and 4 were precoated in the microplate wells. The samples were treated and diluted 50-fold before adding them to the treatment plate and the overall dilution was 100-fold prior to the addition in the NS1 Ab detection well. The secondary Ab used was anti-mouse IgG (HRP-tagged) (abcam, cat# ab97040, UK). The OD of the samples was compared to the OD of anti-dengue virus NS1 glycoprotein Ab (DN3) to estimate the fold change (abcam, cat# ab41616, UK).

### DNA extraction from mice liver tissue

Approximately 25mg of mice liver tissue in RNAlater (Thermofischer Scientific, MA, USA) was taken and lysed using the lysis buffer provided in Qiagen Blood and Tissue kit (Qiagen, Germany) followed by proteinase K and RNase A (4mg/ml) treatment. DNA extraction was done from the lysate according to the manufacturer’s protocol. This was followed by PCR using DV2-NS1 specific forward and reverse primers. Gel electrophoresis was performed with the PCR product using SYBR safe dye (Invitrogen, MA, USA) in a 1% agarose gel. The agarose gel electrophoresis was carried out for 80 mins at 80V and the gel was examined under a UV-transilluminator in Gel Logic (Carestream, CA, USA).

### RNA extraction from mice liver tissue

Approximately 30 mg of mice liver tissue in RNAlater (ThermoFischer Scientific, MA, USA) was triturated and homogenized using blunt 20-gauge needle and syringe. The lysate was then centrifuged for 3 minutes at 13,300 rpm and the supernatant was aspirated out carefully. RNA extraction was done from the supernatant as per the manufacturer’s protocol using Qiagen RNeasy Mini kit (Qiagen, Germany). This was followed by DNase treatment as per the manufacturer’s protocol provided in the Ambion TURBO DNA-free kit (ThermoFischer Scientific, MA, USA). RT-PCR was performed with the elute using DV2 NS1 specific forward and reverse primers stated earlier. The PCR product was subjected to gel electrophoresis in 1% agarose gel with SYBR safe dye (Invitrogen, MA, USA). Agarose gel electrophoresis was done at 80V for 80 mins and the gel was observed in Gel Logic (Carestream, CA, USA) under UV transilluminator.

### CIC C1q ELISA

CIC-C1q levels in mice serum samples from both the experiments were quantified using CIC C1q ELISA kit (DRG International, Cat no: EIA-3169, NJ, USA) according to the manufacturer’s protocol with a few modifications. The principle is based on the interaction between C1q-bound ICs and C1q immobilized on a microplate. The samples were diluted 50-fold before adding them to the microplate. The enzyme conjugate used was 10000-fold diluted anti-mouse IgG (HRP-tagged) (abcam, cat# ab97040). The OD of the samples were measured at 450nm against blank and the IgG concentration of each sample was calculated by interpolating the respective OD values on the standard curve.

### Tissue histopathology

Tissues excised from mice were fixed in 10% formalin (Sigma-Aldrich, MA, USA), dehydrated in 70% to 100% increasing concentrations of ethanol and embedded in paraffin wax (Sigma-Aldrich, MA, USA) in paraffin embedder (HistoCore Arcadia H, LEICA, Germany). Paraffin blocks were sectioned at 5µm using a microtome (HistoCore MULTICUT, LEICA, Germany) and tissue sections were mounted on glass slides coated with Mayer’s albumin (Sigma-Aldrich, USA) and dried overnight. The sections were deparaffinised with xylene and rehydrated with 70% ethanol and water. The rehydrated sections were stained using 90% hematoxylin and 2% eosin (Sigma-Aldrich, USA) for histological observations. The stained sections were mounted using DPX mounting media (Sigma-Aldrich, USA) and observed under bright field microscope (Leica, Germany) at 40x magnification.

### Statistical analysis

Data were analysed using GraphPad Prism 9.0 (GraphPad Software, USA) and reported as mean□±□standard error of mean (SEM). Significant differences between groups were analysed using one-way ANOVA or Student’s t-test and are designated as follows: \**p*□<□0.05, \*\**p*□<□0.01, \*\*\**p*□<□0.001 and \*\*\*\**p*□<□0.0001 relative to the uninfected control groups.

## Supporting information

ARRIVE

## Resource availability

### Lead contact

Request for further information and resources should be directed to and will be fulfilled by the lead contact, Subhajit Biswas.

### Material availability

This study did not generate new unique reagents.

## Acknowledgements

Authors acknowledges the support received from the Directors of CSIR-IICB and NIPER Kolkata, for providing infrastructural support and funding. The project was funded by CSIR-IICB institutional grant to S.B. (MLP-118). S.B. also acknowledges AcSIR for support. S.S. acknowledges the support received from the Director, CHINTA, TCG CREST. C.D. also acknowledges the support of CSIR for her CSIR Research Fellowship.

## Supplementary information

Document S1. Figures S1-S4, Table S1 and S2.

Document S2 / ARRIVE guidelines.

## References

1. G. N. Malavige, G. Ogg, Pathogenesis of severe dengue infection. Ceylon Med J 57, 97–100 (2012).

2. T. P. Htun, Z. Xiong, J. Pang, Clinical signs and symptoms associated with WHO severe dengue classification: a systematic review and meta-analysis. Emerg Microbes Infect 10, 1116–1128 (2021).

3. C. F. Yung, et al., Dengue serotype-specific differences in clinical manifestation, laboratory parameters and risk of severe disease in adults, Singapore. American Journal of Tropical Medicine and Hygiene 92, 999–1005 (2015).

4. Dengue-Global situation. Available at: https://www.who.int/emergencies/disease-outbreak-news/item/2023-DON498 [Accessed 5 September 2025].

5. A. Tayal, S. K. Kabra, R. Lodha, Management of Dengue: An Updated Review. Indian J Pediatr 90, 168–177 (2023).

6. E. M. Plummer, S. Shresta, Mouse models for dengue vaccines and antivirals. J Immunol Methods 410, 34–38 (2014).

7. M. Aguiar, N. Stollenwerk, S. B. Halstead, The Impact of the Newly Licensed Dengue Vaccine in Endemic Countries. PLoS Negl Trop Dis 10 (2016).

8. S. J. Thomas, Is new dengue vaccine efficacy data a relief or cause for concern? NPJ Vaccines 8 (2023).

9. R. J. Kuhn, et al., Structure of dengue virus: Implications for flavivirus organization, maturation, and fusion. Cell 108, 717–725 (2002).

10. A. Ghosh, et al., Non-structural protein 1 (NS1) variants from dengue virus clinical samples revealed mutations that influence NS1 production and secretion. European Journal of Clinical Microbiology and Infectious Diseases 41, 803–814 (2022).

11. S. Nasar, N. Rashid, S. Iftikhar, Dengue proteins with their role in pathogenesis, and strategies for developing an effective anti-dengue treatment: A review. J Med Virol 92, 941–955 (2020).

12. J. M. Mackenzie, M. K. Jones, P. R. Young, Immunolocalization of the Dengue virus nonstructural glycoprotein NS1 suggests a role in viral RNA replication. Virology 220, 232–240 (1996).

13. P. Bhatt, S. P. Sabeena, M. Varma, G. Arunkumar, Current Understanding of the Pathogenesis of Dengue Virus Infection. Curr Microbiol [Preprint] (2021).

14. H. Puerta-Guardo, D. R. Glasner, E. Harris, Dengue Virus NS1 Disrupts the Endothelial Glycocalyx, Leading to Hyperpermeability. PLoS Pathog 12 (2016).

15. J. H. Amorim, R. P. dos S. Alves, S. B. Boscardin, L. C. de S. Ferreira, The dengue virus non-structural 1 protein: Risks and benefits. Virus Res 181, 53–60 (2014).

16. P. R. Beatty, et al., Dengue virus NS1 triggers endothelial permeability and vascular leak that is prevented by NS1 vaccination. Sci Transl Med 7 (2015).

17. K. L. Carpio, A. D. T. Barrett, Flavivirus ns1 and its potential in vaccine development. Vaccines (Basel) 9 (2021).

18. R. P. dos S. Alves, et al., Protective Immunity to Dengue Virus Induced by DNA Vaccines Encoding Nonstructural Proteins in a Lethal Challenge Immunocompetent Mouse Model. Front Med Technol 2 (2020).

19. G. Silva-Santana, et al., Clinical hematological and biochemical parameters in Swiss, BALB/c, C57BL/6 and B6D2F1 Mus musculus. Animal Model Exp Med 3, 304–315 (2020).

20. M. Wszola, et al., Streptozotocin-induced diabetes in a mouse model (Balb/c) is not an effective model for research on transplantation procedures in the treatment of type 1 diabetes. Biomedicines 9 (2021).

21. S. Yook, et al., Molecularly engineered islet cell clusters for diabetes mellitus treatment. Cell Transplant 21, 1775–1789 (2012).

22. D. R. Glasner, H. Puerta-Guardo, P. R. Beatty, E. Harris, The good, the bad, and the shocking: The multiple roles of dengue virus nonstructural protein 1 in protection and pathogenesis. Annu Rev Virol 5, 227–253 (2018).

23. Kerns EH, Di L, Drug-like properties: Concepts, structure design and methods: From ADME to toxicity optimization. Academic Press (2008), San Diego, pp 497–510.

24. A. Al Shoyaib, S. R. Archie, V. T. Karamyan, Intraperitoneal Route of Drug Administration: Should it Be Used in Experimental Animal Studies? Pharm Res 37 (2020).

25. S. M. Costa, et al., Protection against dengue type 2 virus induced in mice immunized with a DNA plasmid encoding the non-structural 1 (NS1) gene fused to the tissue plasminogen activator signal sequence. Vaccine 24, 195–205 (2006).

26. S. M. Costa, et al., DNA vaccines against dengue virus based on the ns1 gene: The influence of different signal sequences on the protein expression and its correlation to the immune response elicited in mice. Virology 358, 413–423 (2007).

27. G. Mellado-Sánchez, et al., A plasmid encoding parts of the dengue virus E and NS1 proteins induces an immune response in a mouse model. Arch Virol 155, 847–856 (2010).

28. P. B. A. Pinto, et al., T cell responses induced by DNA vaccines based on the DENV2 e and NS1 proteins in mice: Importance in protection and immunodominant epitope identification. Front Immunol 10 (2019).

29. K. A. Fontaine, E. L. Sanchez, R. Camarda, M. Lagunoff, Dengue Virus Induces and Requires Glycolysis for Optimal Replication. J Virol 89, 2358–2366 (2015).

30. S. C. Weng, P. N. Tsao, S. H. Shiao, Blood glucose promotes dengue virus infection in the mosquito Aedes aegypti. Parasit Vectors 14, 1–9 (2021).

31. K. Z. Latt, et al., Diabetes mellitus as a prognostic factor for dengue severity: Retrospective study from Hospital for Tropical Diseases, Bangkok. Clin Infect Pract 7–8, 100028 (2020).

32. S. Raj, R. G. V, “International Journal of Advanced Research and Review PROGNOSTIC SIGNIFICANCE OF DIABETES IN DENGUE FEVER WITH POLYSEROSITIS” (2017).

33. R. Singh, et al., Study on dengue severity in diabetic and non-diabetic population of tertiary care hospital by assessing inflammatory indicators. Annals of Medicine and Surgery 82 (2022).

34. C. Dalugama, I. B. Gawarammana, Dengue hemorrhagic fever complicated with transient diabetic ketoacidosis: A case report. J Med Case Rep 11, 10–12 (2017).

35. M. Aamir, et al., Newly diagnosed diabetes mellitus in patients with dengue fever admitted in teaching hospital of Lahore. Pakistan Journal of Medical and Health Sciences 9, 99–101 (2015).

36. S.R. Sudulagunta, M.B. Sodalagunta, S.K. Kothandapani, M.A. Sham, Acute pancreatitis and diabetes mellitus caused by dengue. American Journal of Current Medicine 1:1–10 (2015).

37. M. A. Flor, J. V. Andrade, J. A. Bucaram, Acute Pancreatitis Secondary to Dengue Fever: An Uncommon Presentation of a Common Endemic Illness. Case Rep Infect Dis 2022, 1–7 (2022).

38. X. Liu, et al., Dengue virus is involved in insulin resistance via the downregulation of IRS-1 by inducing TNF-α secretion. Biochim Biophys Acta Mol Basis Dis 1868 (2022).

39. C. T. Wu, et al., SARS-CoV-2 infects human pancreatic β cells and elicits β cell impairment. Cell Metab 33, 1565–1576.e5 (2021).

40. J. A. Müller, et al., SARS-CoV-2 infects and replicates in cells of the human endocrine and exocrine pancreas. Nat Metab 3, 149–165 (2021).

41. K. Mine, S. Nagafuchi, H. Mori, H. Takahashi, K. Anzai, Sars-cov-2 infection and pancreatic β cell failure. Biology (Basel) 11 (2022).

42. K. Khunti, et al., Covid-19, hyperglycemia, and new-onset diabetes. Diabetes Care 44, 2645–2655 (2021).

43. E. K. Kendall, V. R. Olaker, D. C. Kaelber, R. Xu, P. B. Davis, Association of SARS-CoV-2 Infection with New-Onset Type 1 Diabetes among Pediatric Patients from 2020 to 2021. JAMA Netw Open 5, E2233014 (2022).

44. A. H. M. N. Nabi, A. Ebihara, H. U. Shekhar, Impacts of SARS-CoV-2 on diabetes mellitus: A pre and post pandemic evaluation. World J Virol 12, 151–171 (2023).

45. D. Allonso, et al., Dengue Virus NS1 Protein Modulates Cellular Energy Metabolism by Increasing Glyceraldehyde-3-Phosphate Dehydrogenase Activity. J Virol 89, 11871–11883 (2015).

46. C. Chumchanchira, et al., Glycolysis is reduced in dengue virus 2 infected liver cells. Sci Rep 14, 1–14 (2024).

47. E. M. Silva, et al., Dengue virus nonstructural 3 protein interacts directly with human glyceraldehyde-3-phosphate dehydrogenase (GAPDH) and reduces its glycolytic activity. Sci Rep 9, 1–19 (2019).

48. M. Kanwar, R. A. Kowluru, Role of glyceraldehyde 3-phosphate dehydrogenase in the development and progression of diabetic retinopathy. Diabetes 58, 227–234 (2009).

49. B. Giri, et al., Chronic hyperglycemia mediated physiological alteration and metabolic distortion leads to organ dysfunction, infection, cancer progression and other pathophysiological consequences: An update on glucose toxicity. Biomedicine and Pharmacotherapy 107, 306–328 (2018).

50. M. M. Ngwe Tun, Y. Muta, S. Inoue, K. Morita, Persistence of Neutralizing Antibody Against Dengue Virus 2 after 70 Years from Infection in Nagasaki. Biores Open Access 5, 188–191 (2016).

51. J. M. Carr, S. Cabezas-Falcon, J. G. Dubowsky, J. Hulme-Jones, D. L. Gordon, Dengue virus and the complement alternative pathway. FEBS Lett 594, 2543–2555 (2020).

52. R. Kraivong, N. Punyadee, M. K. Liszewski, J. P. Atkinson, P. Avirutnan, Dengue and the lectin pathway of the complement system. Viruses 13 (2021).

53. J. N. Conde, E. M. Silva, A. S. Barbosa, R. Mohana-Borges, The complement system in flavivirus infections. Front Microbiol 8 (2017).

54. P. Avirutnan, et al., Vascular leakage in severe dengue virus infections: A potential role for the nonstructural viral protein NS1 and complement. Journal of Infectious Diseases 193, 1078–1088 (2006).

55. P. Koraka, et al., Detection of immune-complex-dissociated nonstructural-1 antigen in patients with acute dengue virus infections. J Clin Microbiol 41, 4154–4159 (2003).

56. B. N. B. S. F. P. T. P. Boonpucknavig V, Glomerular changes in dengue hemorrhagic fever - PubMed. (1976). Available at: https://pubmed.ncbi.nlm.nih.gov/786215/ [Accessed 5 September 2025].

57. J. F. Picollo Oliveira, E. A. Burdmann, Dengue-associated acute kidney injury. Clin Kidney J 8, 681–685 (2015).

58. A. A. H. Kadhum, et al., Dengue fever-associated glomerulonephritis; an updated narrative mini-review. J Nephropharmacol 14, e12711–e12711 (2024).

59. D. A. Muller, et al., Kinetics of NS1 and anti-NS1 IgG following dengue infection reveals likely early formation of immune complexes in secondary infected patients. Sci Rep 15 (2025).

60. S. M. A. Toloza, S. E. Agüero, Arthritis associated with alphavirus infections: Dengue and Zika. Infections and the Rheumatic Diseases 125–142 (2019). 10.1007/978-3-030-23311-2_12.

61. L. I. Zambrano, et al., Assessment of Post-Dengue Rheumatic Symptoms Using the WOMAC and DAS-28 Questionnaires in a Honduran Population after a Four-Month Follow-Up. Trop Med Infect Dis 7 (2022).

62. W. D. Jayamali, H. M. M. T. B. Herath, A. Kulatunga, A young female presenting with unilateral sacroiliitis following dengue virus infection: A case report. J Med Case Rep 11 (2017).

63. S. Wati, et al., Dengue Virus Infection Induces Upregulation of GRP78, Which Acts To Chaperone Viral Antigen Production. J Virol 83, 12871 (2009).

64. S. Jindadamrongwech, C. Thepparit, D. R. Smith, Identification of GRP 78 (BiP) as a liver cell expressed receptor element for dengue virus serotype 2. Arch Virol 149, 915– 927 (2004).

65. M. O. Aksoy, et al., Secretion of the endoplasmic reticulum stress protein, GRP78, into the BALF is increased in cigarette smokers. Respir Res 18 (2017).

66. M. Gonzalez-Gronow, S. V. Pizzo, Physiological Roles of the Autoantibodies to the 78-Kilodalton Glucose-Regulated Protein (GRP78) in Cancer and Autoimmune Diseases. Biomedicines 10 (2022).

67. S. Denolly, et al., Dengue virus NS1 secretion is regulated via importin-subunit β1 controlling expression of the chaperone GRp78 and targeted by the clinical drug ivermectin. mBio 14 (2023).

68. D. Rondas, et al., Citrullinated glucose-regulated protein 78 is an autoantigen in type 1 diabetes. Diabetes 64, 573–586 (2015).

69. M. Buitinga, et al., Inflammation-induced citrullinated glucose-regulated protein 78 elicits immune responses in human type 1 diabetes. Diabetes 67, 2337–2348 (2018).

70. J. Girona, et al., The circulating GRP78/BIP is a marker of metabolic diseases and atherosclerosis: Bringing endoplasmic reticulum stress into the clinical scenario. J Clin Med 8 (2019).

71. S. H. Back, R. J. Kaufman, Endoplasmic reticulum stress and type 2 diabetes. Annu Rev Biochem 81, 767–793 (2012).

72. C. F. Lin, et al., Generation of IgM anti-platelet autoantibody in dengue patients. J Med Virol 63, 143–149 (2001).

73. S. W. Wan, et al., Autoimmunity in dengue pathogenesis. Journal of the Formosan Medical Association 112, 3–11 (2013).

74. H. J. Cheng, et al., Correlation between serum levels of anti-endothelial cell autoantigen and anti-dengue virus nonstructural protein 1 antibodies in dengue patients. American Journal of Tropical Medicine and Hygiene 92, 989–995 (2015).

75. H. T. M. Vo, et al., Autoantibody profiling in plasma of dengue virus–infected individuals. Pathogens 9, 1–14 (2020).

76. C. M. Filippi, M. G. Von Herrath, Viral trigger for type 1 diabetes: Pros and cons. Diabetes 57, 2863–2871 (2008).

77. R. E. Lloyd, M. Tamhankar, Å. Lernmark, Enteroviruses and Type 1 Diabetes: Multiple Mechanisms and Factors? Annu Rev Med 73, 483–499 (2022).

78. P. De Candia, et al., Type 2 diabetes: How much of an autoimmune disease? Front Endocrinol (Lausanne) 10 (2019).

79. T. Lung, et al., The utility of complement assays in clinical immunology: A comprehensive review. J Autoimmun 95, 191–200 (2018).

80. W. Leowattana, T. Leowattana, Dengue hemorrhagic fever and the liver. World J Hepatol 13, 1968–1976 (2021).

81. A. M. Swamy, P. Y. Mahesh, S. T. Rajashekar, Liver function in dengue and its correlation with disease severity: a retrospective cross-sectional observational study in a tertiary care center in Coastal India. Pan African Medical Journal 40 (2021).

82. C. F. Lin, et al., Liver injury caused by antibodies against dengue virus nonstructural protein 1 in a murine model. Laboratory Investigation 88, 1079–1089 (2008).

83. S. Pahari, et al., Spontaneous splenic hematoma secondary to dengue infection: a rare case report. Ann Med Surg (Lond) 85, 1030–1033 (2023).

84. S. Dronamraju, et al., Splenic artery embolization in subcapsular splenic hematoma secondary to dengue hemorrhagic fever. J Glob Infect Dis 13, 145–147 (2021).

85. K. C. Wei, et al., Major acute cardiovascular events after dengue infection–A population-based observational study. PLoS Negl Trop Dis 16 (2022).

86. S. Yacoub, H. Wertheim, C. P. Simmons, G. Screaton, B. Wills, Cardiovascular manifestations of the emerging dengue pandemic. Nat Rev Cardiol 11, 335–345 (2014).

87. S. Sakinah, et al., Repeated infections of dengue (serotype DENV-2) in lung cells of BALB/c mice lead to severe histopathological consequences. Pathog Glob Health 112, 259–267 (2018).

88. T. F. Póvoa, et al., The Pathology of Severe Dengue in Multiple Organs of Human Fatal Cases: Histopathology, Ultrastructure and Virus Replication. PLoS One 9, 83386 (2014).

89. H.-J. Chen, et al., Transfer of IgG from Long COVID patients induces symptomology in mice. bioRxiv 2024.05.30.596590 (2024). 10.1101/2024.05.30.596590.

90. K. Santos Guedes de Sa, et al., A causal link between autoantibodies and neurological symptoms in long COVID. medRxiv (2024). 10.1101/2024.06.18.24309100,.

91. F. J. Carod-Artal, O. Wichmann, J. Farrar, J. Gascón, Neurological complications of dengue virus infection. Lancet Neurol 12, 906–919 (2013).

92. G. H. Li, Z. J. Ning, Y. M. Liu, X. H. Li, Neurological manifestations of dengue infection. Front Cell Infect Microbiol 7 (2017).

93. S. R. K. Hokkanen, et al., Hippocampal sclerosis, hippocampal neuron loss patterns and TDP-43 in the aged population. Brain Pathology 28, 548–559 (2018).

94. J. Ding, et al., In-host modeling of dengue virus and non-structural protein 1 and the effects of ivermectin in patients with acute dengue fever. CPT Pharmacometrics Syst Pharmacol 13, 2196–2209 (2024).

95. Y. Suputtamongkol, et al., Ivermectin Accelerates Circulating Nonstructural Protein 1 (NS1) Clearance in Adult Dengue Patients: A Combined Phase 2/3 Randomized Double-blinded Placebo Controlled Trial. Clinical Infectious Diseases 72, E586–E593 (2021).

96. N. Kaewjiw, et al., Domperidone inhibits dengue virus infection by targeting the viral envelope protein and nonstructural protein 1. Sci Rep 15 (2025).

97. H. Nath, A. Ghosh, K. Basu, A. De, S. Biswas, Dengue virus clinical isolates sustain viability of infected hepatic cells by counteracting apoptosis-mediated DNA breakage. bioRxiv 2020.06.19.162479 (2021). 10.1101/2020.06.19.162479.

98. R. Raut, et al., Dengue type 1 viruses circulating in humans are highly infectious and poorly neutralized by human antibodies. Proc Natl Acad Sci U S A 116, 227–232 (2019).

99. M. A. Kalas, L. Chavez, M. Leon, P. T. Taweesedt, S. Surani, Abnormal liver enzymes: A review for clinicians. World J Hepatol 13, 1688–1698 (2021).

100. H. Nath, et al., COVID-19 serum can be cross-reactive and neutralizing against the dengue virus, as observed by the dengue virus neutralization test. International Journal of Infectious Diseases 122, 576–584 (2022).

