## Supplementary material for "Dengue NS1 Antibodies drive Immune Complex Formation, Hyperglycaemia and systemic pathology in a murine NS1 plasmid challenge model": Fig. S1

**Short title- “Dengue NS1 Immunopathology in a murine model”**

Chiroshri Dutta^1,6^ Debrupa Dutta^2,6^ Soumi Sukla,^2,3**^ Subhajit Biswas^1,4,5*^

^1^Infectious Diseases and Immunology Division, CSIR-Indian Institute of Chemical Biology (IICB), Kolkata, West Bengal, India

^2^Department of Pharmacology and Toxicology, National Institute of Pharmaceutical Education and Research, Kolkata, West Bengal, India

^3^Present address: CHINTA, TCG Centres for Research and Education in Science and Technology, Kolkata, West Bengal, India

^4^Academy of Scientific and Innovative Research (AcSIR), Ghaziabad, India

^5^Lead contact

^6^These authors contributed equally.

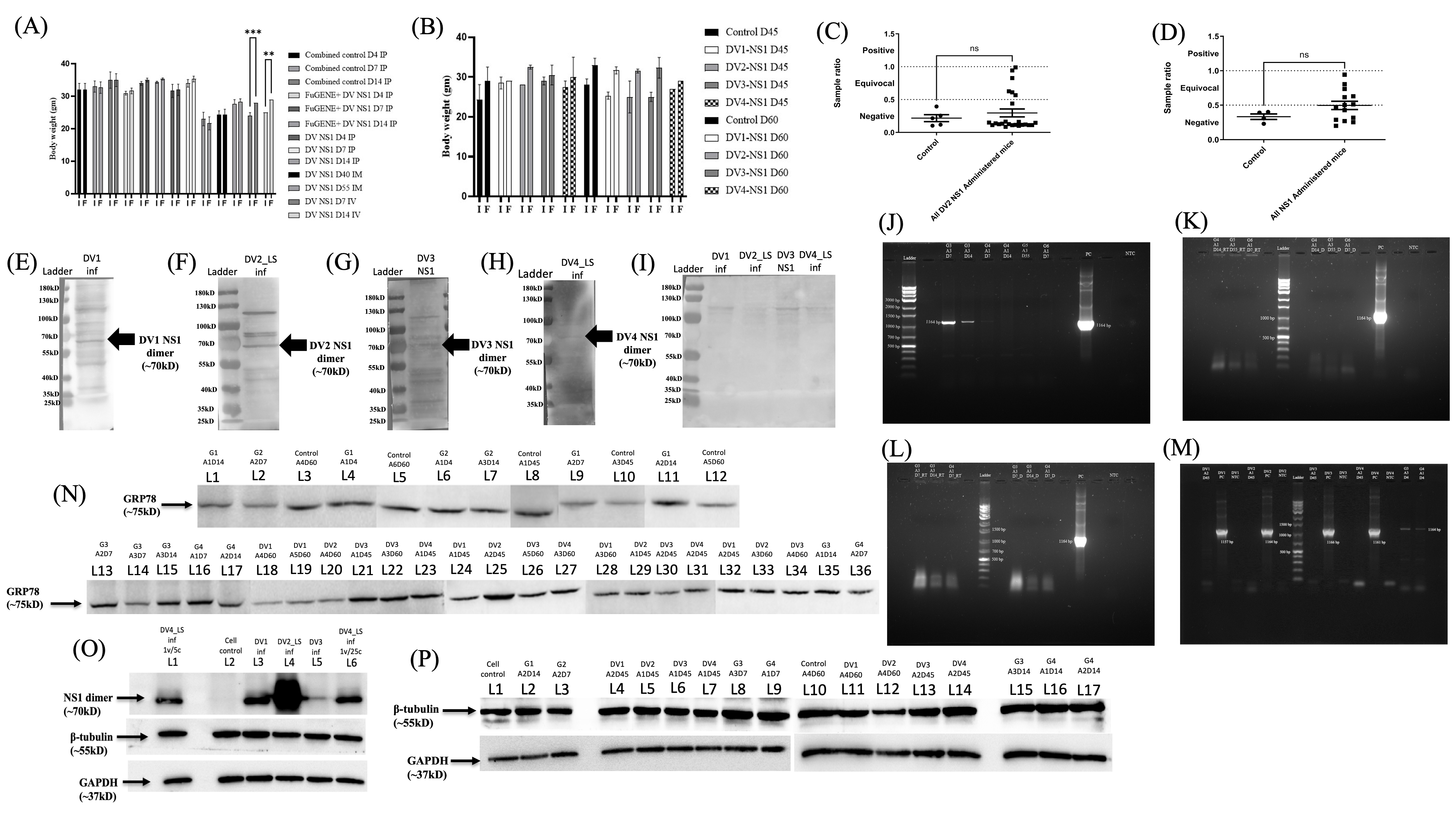

**Fig S1:** **Characterization of body weight changes, NS1 antigen/antibody detection, NS1 DNA/RNA presence, GRP78 level in DV NS1 plasmid-administered mice and GAPDH expression in *in-vitro* models. (A-B)** Body weight analysis of DV NS1 administered mice before the initial dosing (I) and during sacrifice (F). **(A)** Changes in body weight (g) of mice administered with DV2 NS1 plasmid via IP, IM, and IV routes at various time points p.i. **(B)** Body weight changes at sacrifice in mice administered with NS1 plasmids from different DV serotypes (DV1–4) via IP route at 45 and 60 dpi. Data represented as mean ± SEM; **p<0.01, ***p<0.001; D = days post DV NS1 administration. **(C-D) Detection of NS1 antigen in mice sera.** **(C)** ELISA-based quantification of DV2 NS1 antigen in sera of mice administered via IP, IM, and IV routes compared to controls. **(D)** DV NS1 antigen levels in mice administered with NS1 plasmids from DV1–4 serotypes. Data shown as mean ± SEM; OD >1 considered NS1 Ag positive; ns = non-significant. **(E-I)** **Western blot detection of NS1 antibodies** in sera of mice administered with DV1–4 NS1 plasmids and mock-immunized. Cell lysates from Huh7 cells infected with DV1, DV2, DV4 or transfected with DV3 NS1 plasmid were probed with corresponding NS1 Ab-positive or control mouse sera to confirm NS1-specific antibody production.

**(J–M)** **Agarose gel analysis of liver DNA/RNA for presence of NS1 gene.**

**(J)** Gel electrophoresis of DNA extracted from DV2 NS1 Ab-positive mice liver showing 1164 bp DV2 NS1 amplicon.

**(K-L)** Agarose gel image of RNA and its cDNA followed by PCR with DV2 NS1 primers. **(M)** Agarose gel image of DNA extracted from liver of mice administered with different serotypes of DV NS1 plasmid.

Sample nomenclature in J-M: PC = positive control; NTC = no template control; ladder = molecular marker; RT = reverse transcription and D = direct PCR from RNA after DNase treatment.

**(N)** WB of GRP78 expression in serums of control and NS1 plasmid DNA inoculated mice. **(O-P)** WB of **GAPDH protein expression in Huh7 cells. (O)** WB showing GAPDH expression in Huh7 cells infected with DV1–4 serotypes for 96 h. β-tubulin used as internal loading control; NS1 Ag detection confirmed active infection. **(P)** WB of GAPDH expression in Huh7 cells treated with NS1 antibodies for 96 h.

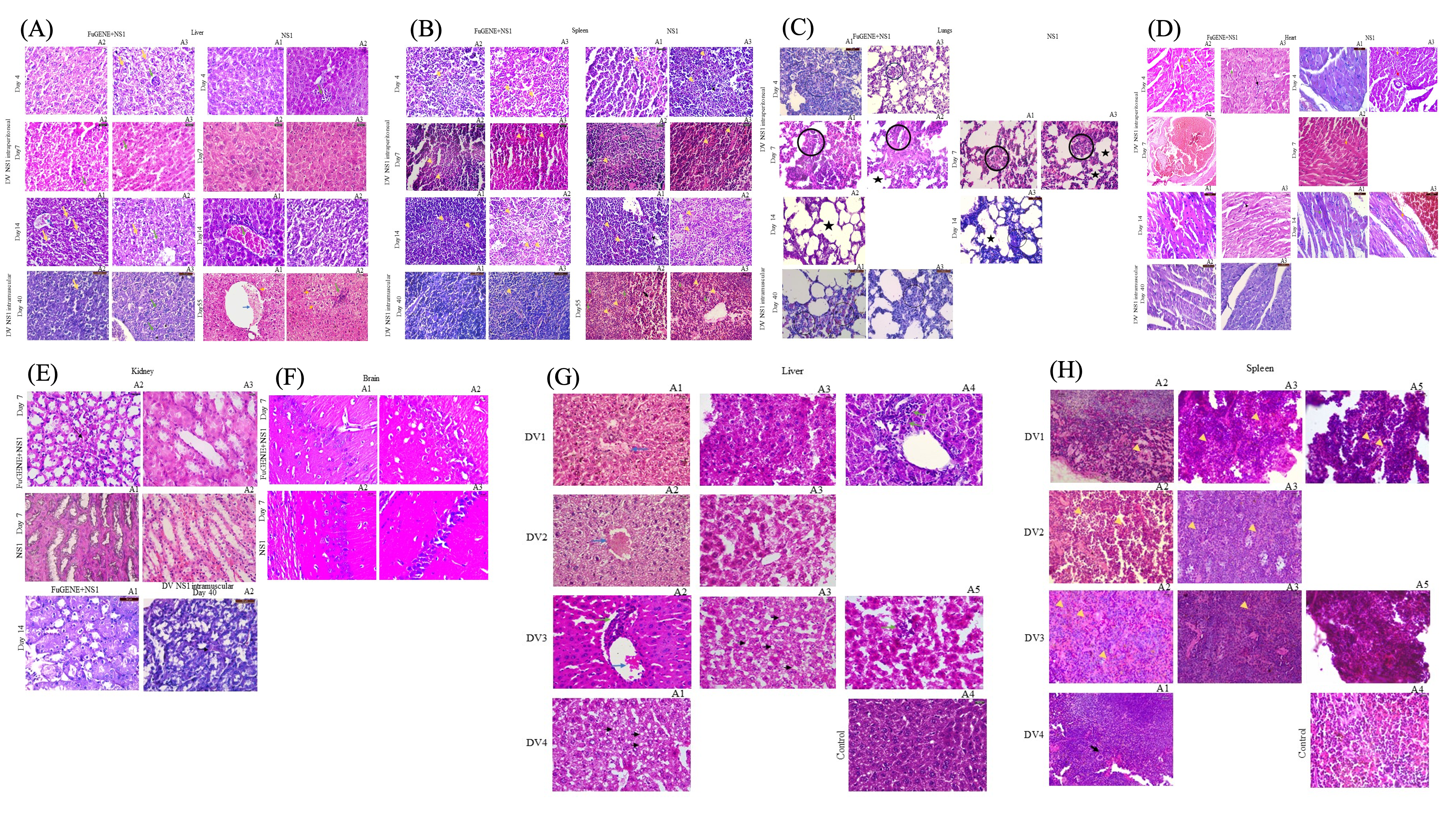

**Fig S2: Histopathological alterations in various organs of DV NS1 plasmid-administered mice across different serotypes, routes, and time points. (A)** Liver sections from DV2 NS1 plasmid-administered mice via IP, IM, and IV routes show progressive pathology including ballooning degeneration of hepatocytes (yellow arrow), mononuclear infiltration (green arrow), and central vein congestion (blue arrow). **(B)** Spleen sections from the same mice reveal severe lymphocyte depletion in the white pulp (yellow arrow), presence of apoptotic bodies (green arrow), and appearance of macrocytic megakaryocytes (black arrow). **(C)** Lung sections from IP and IM groups display structural lung damage in the form of alveolar atelectasis (black circle) and fusion (star). **(D)** Heart tissue shows interstitial edema (yellow arrow), perivascular fibrosis (green arrow), detachment of arteriole from adjacent epithelium (red arrow), and mononuclear infiltration (black arrow). **(E)** Kidney sections demonstrate hyaline cast deposition in renal tubules (black arrow). **(F)** Brain sections show neuronal loss in the CA1 region of the hippocampus. **(G)** Liver sections of mice administered with DV1–4 NS1 plasmids via IP route show consistent central vein congestion (blue arrow), mononuclear infiltration (green arrow), and steatosis (black arrow). **(H)** Spleen sections from these serotype-specific NS1 groups display white pulp lymphocyte depletion (yellow arrow) and increased megakaryocyte presence (black arrow), especially at later time points. All tissues were stained with hematoxylin and eosin and visualized at 40X magnification.

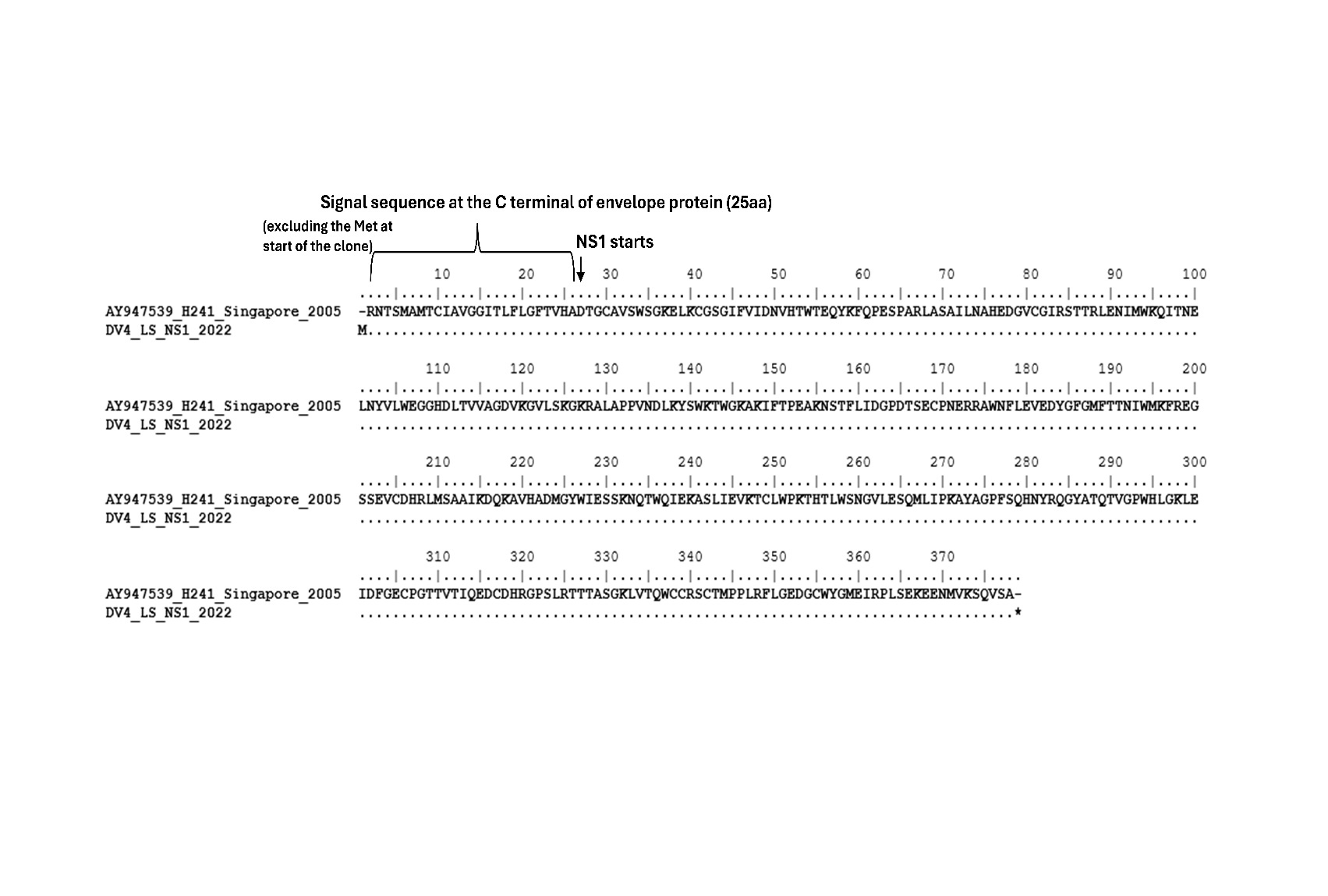

**Fig S3: Sequence alignment of DV4_LS NS1 amino acid.** The sequence alignments of NS1 amino acid of DV4 laboratory strain sequenced from recombinant plasmid DNA with the existing H241 strain already present in GenBank (Accession Number- AY947539).

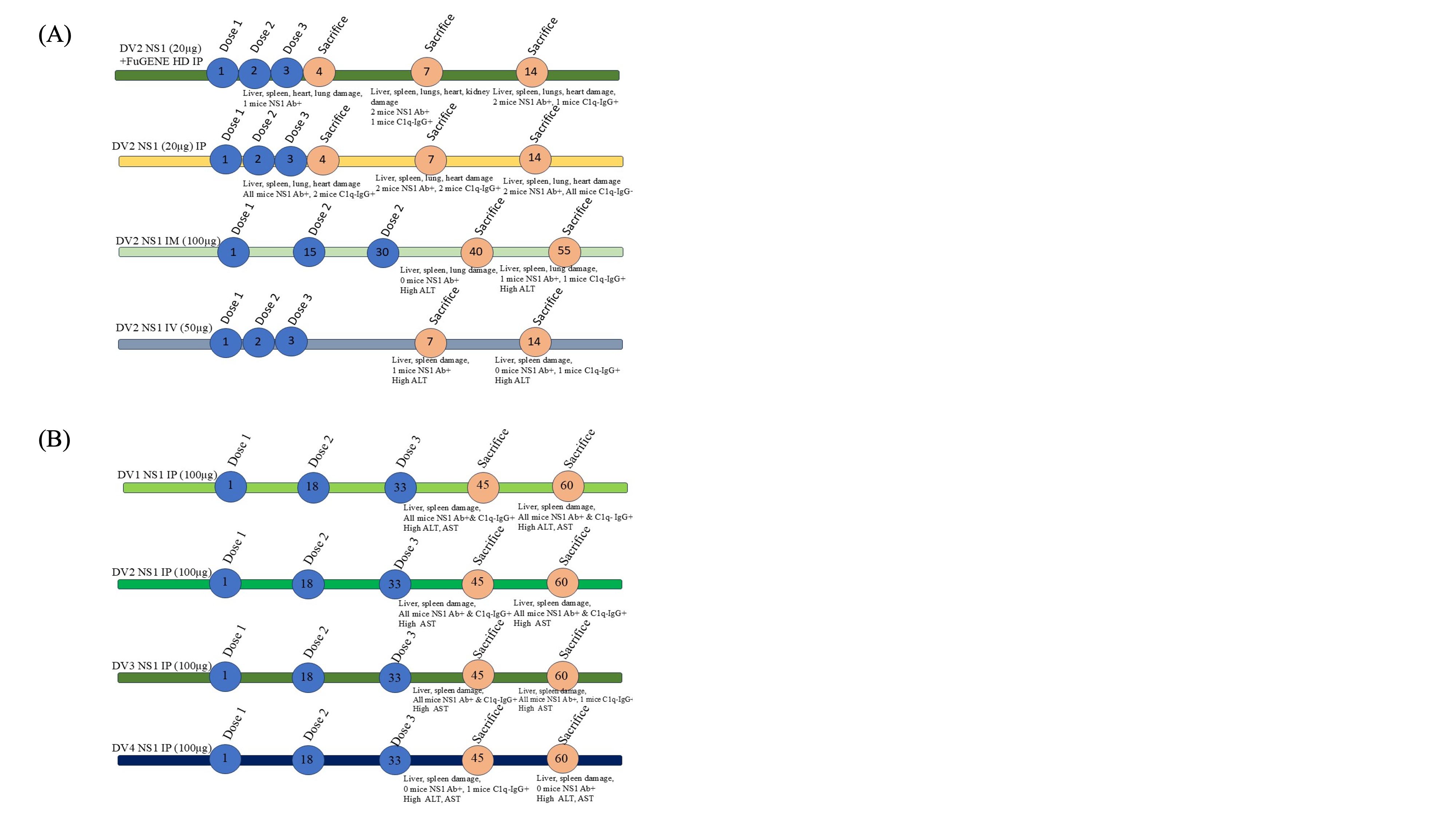

**Fig S4: Schematic representation of experimental design for DV NS1 pathogenesis studies in mice. (A*)*** Experimental Set 1: Mice were administered with DV2 NS1 plasmid via three different routes—intraperitoneal (IP, with or without FuGENE), intramuscular (IM), and intravenous (IV). Animals were sacrificed at multiple timepoints post-administration to assess seroconversion, tissue-specific pathology, molecular changes, and biochemical parameters. **(B)** Experimental Set 2: Mice received 100 µg of NS1-expressing plasmids from all four dengue virus serotypes (DV1–DV4) via IP route in a 3-dose schedule at regular intervals. Post-treatment, mice were evaluated at various timepoints for serotype-specific pathological alterations, immune responses, and tissue damage.

**Table S1: List of different serotype specific dengue virus NS1 cloning primers.**

| **Oligonucleotide primer sequences (5’-3’)** | **Name of primers** | |
| --- | --- | --- |
| CGA AGC TTA GCA TGA GGA RCA CGT CMC TYT CGA TG | | DV1NS1F |
| CGC GGA TCC TTA TGC AGA GAC CAT TGA CCT GAC | | DV1NS1R |
| CGC GCT AGC GCC ATG AAT TCA CGY AGC ACC TC | | DV2NS1F |
| CGC CTC GAG TTA RGC TGT RAC CAA AGA ATT G | | DV2NS1R |
| CGG CTA GCA GCA TGG GRT TGA ATT CAA ARA AYA CWT CC | | DV3NS1F |
| CGC GGA TCC TTA BGC TGA GRC TAA AGA CTT TAC C | | DV3NS1R |
| GCA AAG CTT GAC ATG AGA AAC ACC TCA | | DV4NS1F |
| AGT GAA TTC ATC TTA GGC CGA TAC CTG TGA | | DV4NS1R |

**Table S2: Expression of DV NS1 Ab, DV NS1 Ag, serum glucose level and formation of C1q immune complex in NS1 plasmid administered mice serum (G=Group, D=Day of sacrifice, A=Animal, IP= Intraperitoneal, IM= Intramuscular, IV= Intravenous).**

|  | **Animals** | **NS1 Ag Sample**  **ratio** | **NS1 Ag** | **NS1 Ab OD** | **Fold  change** | **NS1 Ab** | **Serum glucose  level (mg/dl)** | **µg Equiv./ml of  aggregates IgG** | **Formation of  immune complex** | **AST** | **ALT** |
| --- | --- | --- | --- | --- | --- | --- | --- | --- | --- | --- | --- |
| **Control IP** | G1 A1 D4 | 0.118 | Negative | 0.132 | 0.078 | Negative | 146 | 12.523 | Negative | 982.170 | 15.002 |
|  | G1 A2 D4 | n.d. | - | 0.156 | 0.092 | Negative | 198 | 9.667 | Negative | 658.243 | 15.873 |
|  | G1 A1 D7 | 0.256 | Negative | 0.091 | 0.054 | Negative | 240 | 10.874 | Negative | 507.151 | 12.889 |
|  | G1 A2 D7 | 0.217 | Negative | 0.223 | 0.131 | Negative | 206 | 6.118 | Negative | 639.725 | 17.460 |
|  | G1 A1 D14 | n.d. | - | 0.116 | 0.068 | Negative | 184 | 13.957 | Negative | 567.211 | 13.976 |
|  | G1 A2 D14 | 0.394 | Negative | 0.174 | 0.102 | Negative | 211 | 10.481 | Negative | 628.797 | 14.286 |
| **FuGENE control IP** | G2 A1 D4 | n.d. | - | 0.068 | 0.04 | Negative | 228 | 8.762 | Negative | 590.871 | 13.915 |
|  | G2 A2 D7 | n.d. | - | 0.265 | 0.156 | Negative | 193 | 13.461 | Negative | 578.131 | 13.251 |
|  | G2 A3 D14 | 0.103 | Negative | 0.342 | **0.201** | Negative- cut off | 220 | 15.364 | Negative | 647.291 | 13.251 |
| **DV NS1+ FuGENE-IP** | G3 A1 D4 | 0.112 | Negative | 0.135 | 0.08 | Negative | 234 | 5.922 | Negative | 871.151 | 14.821 |
|  | G3 A2 D4 | 0.439 | Negative | 0.049 | 0.029 | Negative | 200 | 6.457 | Negative | 885.711 | 13.734 |
|  | G3 A3 D4 | 0.101 | Negative | 0.66 | 0.388 | Positive | 292 | 9.066 | Negative | 771.051 | 10.595 |
|  | G3 A1 D7 | 0.112 | Negative | 0.139 | 0.082 | Negative | 226 | 18.889 | Positive | 594.511 | 15.304 |
|  | G3 A2 D7 | 0.138 | Negative | 1.51 | 0.889 | Positive | 287 | 10.251 | Negative | 281.472 | 17.417 |
|  | G3 A3 D7 | 0.127 | Negative | 1.864 | 1.098 | Positive | 255 | 14.983 | Negative | 383.391 | 17.598 |
|  | G3 A1 D14 | 0.151 | Negative | 0.298 | 0.175 | Negative | 261 | 15.012 | Negative | 680.051 | 16.029 |
|  | G3 A2 D14 | 0.116 | Negative | 0.692 | 0.408 | Positive | 271 | 20.176 | Positive | 811.091 | 16.813 |
|  | G3 A3 D14 | 0.164 | Negative | 1.98 | 1.166 | Positive | 330 | 23.262 | Positive | 494.411 | 17.961 |
| **DV NS1 only-IP** | G4 A1 D4 | 0.085 | Negative | 0.869 | 0.512 | Positive | 251 | 16.219 | Uncertain | 809.271 | 16.632 |
|  | G4 A2 D4 | 0.12 | Negative | 0.58 | 0.342 | Positive | 226 | 45.546 | Positive | 958.510 | 13.312 |
|  | G4 A3 D4 | 0.111 | Negative | 0.67 | 0.395 | Positive | 308 | 46.801 | Positive | 674.591 | 13.674 |
|  | G4 A1 D7 | 0.125 | Negative | 1.46 | 0.86 | Positive | 282 | 27.401 | Positive | 918.471 | 10.051 |
|  | G4 A2 D7 | 0.116 | Negative | 0.196 | 0.115 | Negative | 342 | 11.302 | Negative | 467.111 | 9.931 |
|  | G4 A3 D7 | 0.133 | Negative | 0.541 | 0.318 | Positive | 224 | 23.468 | Positive | 652.751 | 17.115 |
|  | G4 A1 D14 | 0.112 | Negative | 2.304 | 1.357 | Positive | 231 | 26.871 | Positive | 425.251 | 9.629 |
|  | G4 A2 D14 | 0.16 | Negative | 1.193 | 0.703 | Positive | 276 | 27.854 | Positive | 592.691 | 16.270 |
|  | G4 A3 D14 | 0.147 | Negative | 0.194 | 0.114 | Negative | 172 | 28.215 | Positive | 772.871 | 5.584 |
| **DV NS1 only-IM** | G5 A1 D40 | 0.827 | Equivocal | 0.062 | 0.037 | Negative | 251 | 8.063 | Negative | 854.771 | 22.549 |
|  | G5 A2 D40 | n.d. | - | 0.035 | 0.021 | Negative | 214 | 5.134 | Negative | 789.251 | 21.100 |
|  | G5 A3 D40 | 0.626 | Equivocal | 0.054 | 0.032 | Negative | 186 | 4.15 | Negative | 787.431 | 18.987 |
|  | G5 A1 D55 | n.d. | - | 0.05 | 0.029 | Negative | 191 | 9.67 | Negative | 836.571 | 43.983 |
|  | G5 A2 D55 | n.d. | - | 0.056 | 0.033 | Negative | 237 | 7.281 | Negative | 734.651 | 17.598 |
|  | G5 A3 D55 | 0.609 | Equivocal | 1.583 | 0.933 | Positive | 322 | 65.081 | Positive | 656.391 | 25.991 |
|  | G5 A4 D55 | 0.567 | Equivocal | 0.03 | 0.017 | Negative | 249 | 17.207 | Uncertain | 812.911 | 20.255 |
| **DV NS1 only-IV** | G6 A1 D7 | 0.943 | Equivocal | 1.067 | 0.629 | Positive | 271 | 10.265 | Negative | 681.871 | 23.757 |
|  | G6 A2 D7 | n.d. | - | 0.024 | 0.014 | Negative | 229 | 3.685 | Negative | 685.511 | 22.730 |
|  | G6 A1 D14 | 0.989 | Equivocal | 0.037 | 0.022 | Negative | 360 | 34.547 | Positive | 985.810 | 25.145 |
| **DV1-NS1 IP** | A1 D45 | 0.253 | Negative | 1.239 | 0.73 | Positive | 187 | 27.103 | Positive | 1316.200 | 17.293 |
|  | A2 D45 | 0.428 | Negative | 1.712 | 1.009 | Positive | 158 | 27.628 | Positive | 1042.170 | 39.280 |
|  | A3 D60 | 0.288 | Negative | 0.377 | 0.222 | Positive | 111 | 30.218 | Positive | 1101.960 | 38.632 |
|  | A4 D60 | 0.269 | Negative | 2.572 | 1.515 | Positive | 86 | 28.544 | Positive | 1336.130 | 18.103 |
|  | A5 D60 | 0.616 | Equivocal | 1.414 | 0.833 | Positive | 92 | 36.195 | Positive | 1189.150 | 17.293 |
| **DV2-NS1 IP** | A1 D45 | 0.736 | Equivocal | 2.104 | 1.239 | Positive | 199 | 20.487 | Positive | 1194.130 | 17.901 |
|  | A2 D45 | 0.629 | Equivocal | 1.204 | 0.709 | Positive | 139 | 22.973 | Positive | 1388.440 | 15.997 |
|  | A3 D60 | n.d. | - | 1.193 | 0.703 | Positive | 119 | 22.217 | Positive | 1258.900 | 14.580 |
|  | A4 D60 | 0.804 | Equivocal | 2.276 | 1.341 | Positive | 115 | 35.77 | Positive | 1149.290 | 6.846 |
| **DV3-NS1 IP** | A1 D45 | 0.201 | Negative | 1.653 | 0.973 | Positive | 154 | 44.872 | Positive | 1273.850 | 11.989 |
|  | A2 D45 | 0.512 | Equivocal | 1.662 | 0.979 | Positive | 189 | 23.262 | Positive | 1154.270 | 60.214 |
|  | A3 D60 | n.d. | - | 1.605 | 0.946 | Positive | 102 | 45.015 | Positive | 1209.080 | 14.013 |
|  | A4 D60 | 0.943 | Equivocal | 0.966 | 0.569 | Positive | 203 | 12.325 | Negative | 867.794 | 7.616 |
|  | A5 D60 | 0.502 | Equivocal | 0.584 | 0.344 | Positive | 160 | 34.135 | Positive | 994.841 | 12.434 |
| **DV4-NS1 IP** | A1 D45 | n.d. | - | 0.166 | 0.098 | Negative | 140 | 21.926 | Positive | 780.605 | 11.746 |
|  | A2 D45 | 0.46 | Negative | 0.174 | 0.103 | Negative | 307 | 11.111 | Negative | 1009.790 | 46.650 |
|  | A3 D60 | 0.318 | Negative | 0.182 | 0.107 | Negative | 169 | 13.134 | Negative | 1211.570 | 87.627 |
| **Control D45** | A1 D45 | 0.402 | Negative | 0.089 | 0.052 | Negative | 148 | 14.135 | Negative | 639.726 | 15.002 |
|  | A2 D45 | 0.392 | Negative | 0.138 | 0.081 | Negative | 129 | 3.986 | Negative | 1074.690 | 12.889 |
|  | A3 D45 | n.d. | - | 0.115 | 0.068 | Negative | 85 | 14.942 | Negative | 628.797 | 13.976 |
| **Control D60** | A4 D60 | n.d. | - | 0.077 | 0.045 | Negative | 105 | 13.747 | Negative | 658.244 | 12.698 |
|  | A5 D60 | 0.308 | Negative | 0.21 | 0.124 | Negative | 88 | 8.216 | Negative | 1107.460 | 37.037 |
|  | A6 D60 | 0.23 | Negative | 0.164 | 0.097 | Negative | 120 | 10.288 | Negative | 646.957 | 17.813 |
| Positive control | DN3 |  |  | 1.698 |  |  |  |  |  |  |  |

**Table S3: Fold-change of DV NS1 Ab, DV NS1 Ag, serum glucose level and formation of C1q immune complex in NS1 plasmid administered mice serum along with their respective histo-pathologies. (G=Group, D=Day of sacrifice, A=Animal, IP= Intraperitoneal, IM= Intramuscular, IV= Intravenous).**

| Set 1 Experiment | **Animals** | **Serum glucose   (fold-change w.r.t. mean)** | **NS1 Ab (fold  change w.r.t. positive control)** | **Immune complex formation (fold-change w.r.t. cut-off)** | **AST (fold-change w.r.t. mean)** | **ALT (fold-change w.r.t. mean)** | **Liver** | **Spleen** | **Heart** | **Lungs** | **Kidney** | **Brain** |
| --- | --- | --- | --- | --- | --- | --- | --- | --- | --- | --- | --- | --- |
| Control | G1 A1 D4 | 0.739 | 0.078  (-) | 0.696  (-) | **1.479** | **1.006** | n.d. | n.d. | n.d. | n.d. | n.d. | n.d. |
|  | G1 A2 D4 | **1.003** | 0.092  (-) | 0.537  (-) | 0.992 | **1.064** | n.d. | n.d. | n.d. | n.d. | n.d. | n.d. |
|  | G1 A1 D7 | **1.215** | 0.054  (-) | 0.604  (-) | 0.764 | 0.864 | n.d. | n.d. | n.d. | n.d. | n.d. | n.d. |
|  | G1 A2 D7 | **1.043** | 0.131  (-) | 0.340  (-) | 0.964 | **1.171** | Normal hepatic architecture. | Normal distribution of lymphocytes. | Normal cardiac muscle histology. | Normal alveolar structure. | Normal nephron ultrastructure. | Dense intact pyramidal neurons in the CA1 region. |
|  | G1 A1 D14 | 0.932 | 0.068  (-) | 0.775  (-) | 0.854 | 0.937 | n.d. | n.d. | n.d. | n.d. | n.d. | n.d. |
|  | G1 A2 D14 | **1.068** | 0.102  (-) | 0.582  (-) | 0.947 | 0.958 | n.d. | n.d. | n.d. | n.d. | n.d. | n.d. |
| FuGENE control | G2 A1 D4 | **1.067** | 0.040  (-) | 0.487  (-) | 0.976 | **1.033** | n.d. | n.d. | n.d. | n.d. | n.d. | n.d. |
|  | G2 A2 D7 | 0.903 | 0.156  (-) | 0.748  (-) | 0.955 | 0.984 | Normal hepatic architecture. | Normal distribution of lymphocytes. | Normal cardiac muscle histology. | Normal alveolar structure. | Normal nephron ultrastructure. | Dense intact pyramidal neurons in the CA1 region. |
|  | G2 A3 D14 | **1.030** | 0.201  (-) | 0.854  (-) | **1.069** | 0.984 | n.d. | n.d. | n.d. | n.d. | n.d. | n.d. |
| DV2-NS1+ FuGENE-IP | G3 A1 D4 | **1.095** | 0.080  (-) | 0.329  (-) | **1.439** | **1.100** | Mononuclear infiltration | Severe depletion in lymphocytes from white pulp region. | Interstitial edema & perivascular fibrosis. | Alveolar damage in the form of atelectasis. | n.d. | n.d. |
|  | G3 A2 D4 | 0.936 | 0.029  (-) | 0.359  (-) | **1.463** | **1.019** | Normal hepatic architecture. | Depletion of lymphocytes. | Interstitial edema. | Atelectasis & alveolar fusion. | n.d. | n.d. |
|  | G3 A3 D4 | **1.367** | **0.388**  (+) | 0.504  (-) | **1.274** | 0.786 | Mononuclear infiltration &  ballooning degeneration of hepatocytes. | Depletion of lymphocytes. | Perivascular fibrosis & mononuclear infiltration. | Alveolar damage in the form of atelectasis. | Normal nephron ultrastructure. | n.d. |
|  | G3 A1 D7 | **1.058** | 0.082  (-) | **1.049**  (+) | 0.982 | **1.136** | Mononuclear infiltration &  ballooning of hepatocytes. | Severe depletion in lymphocytes from white pulp region. | n.d. | Alveolar damage in the form of atelectasis. | Atopic degeneration of renal tubules and edema. | Depletion of neurons in CA1 hippocampal region. |
|  | G3 A2 D7 | **1.343** | **0.889**  (+) | 0.569  (-) | 0.465 | **1.293** | Mononuclear infiltration. | Depletion of lymphocytes. | Normal cardiac muscle histology. | Atelectasis & alveolar fusion. | Deposition of hyaline cast. | Depletion of neurons in CA1 hippocampal region. |
|  | G3 A3 D7 | **1.193** | **1.098**  (+) | 0.832  (-) | 0.633 | **1.306** | Mononuclear infiltration. | Severe depletion in lymphocytes from white pulp region. | Interstitial edema. | Atelectasis & alveolar fusion. | Normal nephron ultrastructure. | Diffuse neurons in the CA1 region of brain hippocampus & low neuron count. |
|  | G3 A1 D14 | **1.222** | 0.175  (-) | 0.834  (-) | **1.123** | **1.190** | Central vein congestion & ballooning degeneration of hepatocytes. | Depletion of lymphocytes. | Normal cardiac muscle histology. | n.d. | Normal nephron ultrastructure. | n.d. |
|  | G3 A2 D14 | **1.268** | **0.408**  (+) | **1.121**  (+) | **1.340** | **1.248** | Mononuclear infiltration &  ballooning degeneration of hepatocytes. | Severe depletion in lymphocytes from white pulp region. | Interstitial edema, cardiac degeneration with vascular leakage & arteriolar detachment from peripheral tissues. | Alveolar fusion. | Deposition of hyaline cast. | n.d. |
|  | G3 A3 D14 | **1.544** | **1.166**  (+) | **1.292**  (+) | 0.817 | **1.333** | Maximum ballooning of hepatocytes & portal congestion indicating fibrosis. | Severe depletion in lymphocytes from white pulp region & appearance of macrocytic megakaryocyte. | Mononuclear infiltration. | Alveolar fusion | n.d. | n.d. |
| DV2-NS1 only-IP | G4 A1 D4 | **1.271** | **0.512**  (+) | 0.901  (-/+) | **1.219** | **1.115** | Normal hepatic architecture. | Depletion of lymphocytes. | Normal cardiac muscle histology. | n.d. | Normal nephron ultrastructure. | n.d. |
|  | G4 A2 D4 | **1.144** | **0.342**  (+) | **2.530**  (+) | **1.444** | 0.893 | Mononuclear infiltration. | Severe depletion in lymphocytes from white pulp region. | Interstitial edema & cardiac muscle mononuclear infiltration. | n.d. | n.d. | n.d. |
|  | G4 A3 D4 | **1.559** | **0.395**  (+) | **2.600**  (+) | **1.016** | 0.917 | Presence of fat globules. | Severe depletion in lymphocytes from white pulp region & appearance of macrocytic megakaryocyte. | Interstitial edema & detachment of arteriole from surrounding epithelium. | Atelectasis & alveolar fusion. | n.d. | n.d. |
|  | G4 A1 D7 | **1.428** | **0.860**  (+) | **1.522**  (+) | **1.383** | 0.674 | Mononuclear infiltration. | Severe depletion in lymphocytes from white pulp region & appearance of macrocytic megakaryocyte. | n.d. | Alveolar damage in the form of atelectasis. | Normal nephron ultrastructure. | Diffuse neuron in the CA1 region of brain hippocampus & low neuron count. |
|  | G4 A2 D7 | **1.732** | 0.115  (-) | 0.628  (-) | 0.704 | 0.666 | Normal hepatic architecture. | Depletion of lymphocytes. | Interstitial edema. | Atelectasis | Normal nephron ultrastructure. | Depletion of neurons in CA1 hippocampal region. |
|  | G4 A3 D7 | **1.134** | **0.318**  (+) | **1.304**  (+) | 0.983 | **1.148** | Normal hepatic architecture. | Depletion of lymphocytes. | Cardiac muscle mononuclear infiltration indicating inflammation. | Alveolar damage in the form of atelectasis & alveolar fusion. | Normal nephron ultrastructure. | Depletion of neurons in CA1 hippocampal region. |
|  | G4 A1 D14 | **1.170** | **1.357**  (+) | **1.493**  (+) | 0.641 | 0.646 | Mononuclear infiltration. | Severe depletion in lymphocytes from white pulp region & appearance of macrocytic megakaryocyte. | Perivascular fibrosis. | n.d. | Normal nephron ultrastructure. | n.d. |
|  | G4 A2 D14 | **1.397** | **0.703**  (+) | **1.547**  (+) | 0.893 | **1.091** | Normal hepatic architecture. | Severe depletion in lymphocytes from white pulp region. | Interstitial edema & cardiac muscle mononuclear infiltration. | Alveolar damage in the form of atelectasis. | n.d. | n.d. |
|  | G4 A3 D14 | 0.871 | 0.114  (-) | **1.568**  (+) | **1.164** | 0.374 | Mononuclear infiltration. | Foamy spleen. | Interstitial edema. | Alveolar damage in the form of atelectasis & alveolar fusion. | n.d. | n.d. |
| DV2-NS1 only-IM | G5 A1 D40 | **1.271** | 0.037  (-) | 0.448  (-) | **1.288** | **1.512** | Ballooning degeneration of hepatocytes. | Normal distribution of lymphocytes. | Normal cardiac muscle histology. | Alveolar damage in the form of atelectasis. | Normal nephron ultrastructure. | n.d. |
|  | G5 A2 D40 | **1.084** | 0.021  (-) | 0.285  (-) | **1.189** | **1.415** | Ballooning degeneration of hepatocytes. | Severe depletion in lymphocytes from white pulp region & appearance of macrocytic megakaryocyte. | Normal cardiac muscle histology. | Alveolar damage in the form of atelectasis. | Deposition of hyaline cast. | n.d. |
|  | G5 A3 D40 | 0.942 | 0.032  (-) | 0.231  (-) | **1.186** | **1.273** | Mononuclear infiltration. | Depletion of lymphocytes. | Normal cardiac muscle histology. | Atelectasis | n.d. | n.d. |
|  | G5 A1 D55 | 0.967 | 0.029  (-) | 0.537  (-) | **1.260** | **2.949** | Central vein congestion & ballooning degeneration of hepatocytes. | Severe depletion in lymphocytes from white pulp region & formation of apoptotic bodies. | n.d. | n.d. | n.d. | n.d. |
|  | G5 A2 D55 | **1.200** | 0.033  (-) | 0.405  (-) | **1.107** | **1.180** | Mononuclear infiltration &  ballooning degeneration of hepatocytes. | Severe depletion in lymphocytes, presence of apoptotic bodies & appearance of macrocytic megakaryocyte. | n.d. | n.d. | n.d. | n.d. |
|  | G5 A3 D55 | **1.630** | **0.933**  (+) | **3.616**  (+) | 0.989 | **1.743** | Severe central vein congestion & ballooning degeneration of hepatocytes. | Severe depletion in lymphocytes from white pulp region & formation of apoptotic bodies. | n.d. | n.d. | n.d. | n.d. |
|  | G5 A4 D55 | **1.261** | 0.017  (-) | 0.956  (-/+) | **1.224** | **1.358** | n.d. | n.d. | n.d. | n.d. | n.d. | n.d. |
| DV2-NS1 only-IV | G6 A1 D7 | **1.372** | **0.629**  (+) | 0.570  (-) | **1.027** | **1.593** | Mild hepatic portal vein congestion | n.d. | n.d. | n.d. | n.d. | n.d. |
|  | G6 A2 D7 | **1.159** | 0.014  (-) | 0.205  (-) | **1.033** | **1.524** | n.d. | Severe depletion in lymphocytes from white pulp region & appearance of macrocytic megakaryocyte. | n.d. | n.d. | n.d. | n.d. |
|  | G6 A1 D14 | **1.823** | 0.022  (-) | **1.919**  (+) | **1.485** | **1.686** | Increased hepatic portal vein congestion | Severe depletion in lymphocytes from white pulp region. | n.d. | n.d. | n.d. | n.d. |
| Set 2 Experiment |  |  |  |  |  |  |  |  |  |  |  |  |
| DV1-NS1-IP | A1 D45 | **1.655** | **0.730**  (+) | **1.506**  (+) | **1.661** | 0.948 | Central vein congestion. | Depletion in splenocytes. | n.d. | n.d. | n.d. | n.d. |
|  | A2 D45 | **1.398** | **1.009**  (+) | **1.535**  (+) | **1.315** | **2.154** | Mononuclear infiltration & congestion of central vein. Presence of mild steatosis. | Depletion of lymphocytes. | n.d. | n.d. | n.d. | n.d. |
|  | A3 D60 | 0.982 | **0.222**  (+) | **1.679**  (+) | **1.390** | **2.118** |  | Depletion of lymphocytes. | n.d. | n.d. | n.d. | n.d. |
|  | A4 D60 | 0.761 | **1.515**  (+) | **1.586**  (+) | **1.686** | 0.993 | Mononuclear infiltration. | Depletion in splenocytes. | n.d. | n.d. | n.d. | n.d. |
|  | A5 D60 | 0.814 | **0.833**  (+) | **2.011**  (+) | **1.500** | 0.948 | Ballooning of hepatocytes. | Depletion of lymphocytes. | n.d. | n.d. | n.d. | n.d. |
| DV2-NS1-IP | A1 D45 | **1.761** | **1.239**  (+) | **1.138**  (+) | **1.507** | 0.982 | Central vein congestion & ballooning degeneration of hepatocytes. | Depletion in splenocytes. | n.d. | n.d. | n.d. | n.d. |
|  | A2 D45 | **1.230** | **0.709**  (+) | **1.276**  (+) | **1.752** | 0.877 | Central vein congestion. | Depletion of lymphocytes. | n.d. | n.d. | n.d. | n.d. |
|  | A3 D60 | **1.053** | **0.703**  (+) | **1.234**  (+) | **1.588** | 0.800 |  | Depletion of lymphocytes. | n.d. | n.d. | n.d. | n.d. |
|  | A4 D60 | **1.018** | **1.341**  (+) | **1.987**  (+) | **1.450** | 0.375 | Mononuclear infiltration. | Appearance of multiple megakaryocytes. | n.d. | n.d. | n.d. | n.d. |
|  | A5 D60 | Animal died in the 1^st^ week after the 1^st^ dose. | | | | | | | | | | |
| DV3-NS1-IP | A1 D45 | **1.363** | **0.973**  (+) | **2.493**  (+) | **1.607** | 0.657 | Central vein congestion. | Depletion in splenocytes. | n.d. | n.d. | n.d. | n.d. |
|  | A2 D45 | **1.673** | **0.979**  (+) | **1.292**  (+) | **1.456** | **3.302** | Central vein congestion & mononuclear infiltration. | Depletion of lymphocytes. | n.d. | n.d. | n.d. | n.d. |
|  | A3 D60 | 0.903 | **0.946**  (+) | **2.501**  (+) | **1.525** | 0.768 | Steatosis. | Depletion of lymphocytes. | n.d. | n.d. | n.d. | n.d. |
|  | A4 D60 | **1.796** | **0.569**  (+) | 0.685  (-) | **1.095** | 0.418 | Mononuclear infiltration. | n.d. | n.d. | n.d. | n.d. | n.d. |
|  | A5 D60 | **1.416** | **0.344**  (+) | **1.896**  (+) | **1.255** | 0.682 | Mononuclear infiltration. | n.d. | n.d. | n.d. | n.d. | n.d. |
| DV4-NS1-IP | A1 D45 | **1.239** | 0.098  (-) | **1.218**  (+) | 0.985 | 0.644 | Steatosis. | Appearance of megakaryocytes. | n.d. | n.d. | n.d. | n.d. |
|  | A2 D45 | **2.717** | 0.103  (-) | 0.617  (-) | **1.274** | **2.558** | Central vein congestion. | Depletion in splenocytes & appearance of multiple megakaryocytes. | n.d. | n.d. | n.d. | n.d. |
|  | A3 D60 | **1.496** | 0.107  (-) | 0.730  (-) | **1.529** | **4.805** | Presence of mild steatosis. | Depletion in splenocytes. | n.d. | n.d. | n.d. | n.d. |
|  | A4 D60 | Animal died in the 2^nd^ week after the 1^st^ dose. | | | | | | | | | | |
|  | A5 D60 | Animal died in the 1^st^ week after the 3^rd^ dose. | | | | | | | | | | |
| Vehicle Control | A1 D45 | **1.310** | 0.052  (-) | 0.785  (-) | 0.807 | 0.823 | n.d. | n.d. | n.d. | n.d. | n.d. | n.d. |
|  | A2 D45 | **1.142** | 0.081  (-) | 0.221  (-) | **1.356** | 0.707 | n.d. | n.d. | n.d. | n.d. | n.d. | n.d. |
|  | A3 D45 | 0.752 | 0.068  (-) | 0.830  (-) | 0.793 | 0.766 | n.d. | n.d. | n.d. | n.d. | n.d. | n.d. |
|  | A4 D60 | 0.929 | 0.045  (-) | 0.764  (-) | 0.830 | 0.696 | Normal hepatic architecture. | Normal distribution of lymphocytes. | n.d. | n.d. | n.d. | n.d. |
|  | A5 D60 | 0.779 | 0.124  (-) | 0.456  (-) | **1.397** | **2.031** | n.d. | n.d. | n.d. | n.d. | n.d. | n.d. |
|  | A6 D60 | **1.062** | 0.097 | 0.572 | 0.816 | 0.977 | n.d. | n.d. | n.d. | n.d. | n.d. | n.d. |
| Positive control | DN3 |  | 1.698 |  |  |  |  |  |  |  |  |  |

Mice serum samples having a fold change of >0.201 with respect to the OD of the anti-dengue virus NS1 glycoprotein antibody (DN3) were considered antibody positive. For serum glucose, AST and ALT levels, the fold change values were calculated with respect to the means of the respective control groups. Lastly, for immune complex formation, the fold change was calculated with respect to the cut-off value (i.e.18) already specified in the kit. A fold-change of ≥1 indicates positive result for immune complex formation.
